# Haploid, diploid, and pooled exome capture recapitulate features of biology and paralogy in two non-model tree species

**DOI:** 10.1101/2020.10.07.329961

**Authors:** Brandon Lind, Mengmeng Lu, Dragana Obreht Vidakovic, Pooja Singh, Tom Booker, Sam Yeaman, Sally Aitken

**Author notes:** equal contribution (Brandon Lind, Mengmeng Lu). **Corresponding Author**, Brandon Lind, 2424 Main Mall, 3027 Forest Science Centre, University of British Columbia, Vancouver, BC, Canada.

## Abstract

Despite their suitability for studying evolution, many conifer species have large and repetitive giga-genomes (16-31Gbp) that create hurdles to producing high coverage SNP datasets that captures diversity from across the entirety of the genome. Due in part to multiple ancient whole genome duplication events, gene family expansion and subsequent evolution within *Pinaceae*, false diversity from the misalignment of paralog copies create further challenges in accurately and reproducibly inferring evolutionary history from sequence data. Here, we leverage the cost-saving benefits of pool-seq and exome-capture to discover SNPs in two conifer species, Douglas-fir (*Pseudotsuga menziesii* var. *menziesii* (Mirb.) Franco, *Pinaceae*) and jack pine (*Pinus banksiana* Lamb., *Pinaceae*). We show, using minimal baseline filtering, that allele frequencies estimated from pooled individuals show a strong positive correlation with those estimated by sequencing the same population as individuals (r > 0.948), on par with such comparisons made in model organisms. Further, we highlight the use of haploid megagametophyte tissue in identifying sites that are likely due to misaligned paralogs. Together with additional minor filtering, we show that it is possible to remove many of the loci with large frequency estimate discrepancies between individual and pooled sequencing approaches, improving the correlation further (r > 0.973). Our work addresses bioinformatic challenges in non-model organisms with large and complex genomes, highlights the use of megagametophyte tissue for the identification of paralog sites when sequencing large numbers of populations, and suggests the combination of pool-seq and exome capture to be robust for further evolutionary hypothesis testing in these systems.

## 1 Introduction

Quantifying the spatial structure of neutral and adaptive genetic variation within ecologically and economically important tree species (and close relatives) is fundamental to forecasting and managing their response to changing selection pressures from pests, pathogens, and climate (Aitken et al. 2008; Alberto et al. 2013; Holliday et al. 2017; Janes et al. 2017; Sniezko et al. 2017). Prerequisite to this information is the ability to produce high quality and cost-effective data from which to generate reliable inference. While the life history of many tree species offers some ideal circumstances for studying adaptive evolution at the genetic level (Neale & Savolainen 2004; Neale & Kremer 2011), two ancient whole genome duplication events in progenitors of *Pinaceae* lineages (Li et al. 2015), transposable element dynamics (Morse et al. 2009; Voronova et al. 2017), tandemly arrayed genes (Pavy et al. 2017), subsequent gene duplication (Krutovsky et al. 2004, Casola & Koralewski 2018) and gene family expansion (e.g., Liu et al. 2016) have led to giga-genomes (>16Gb in size) recalcitrant to chromosome-level genome assembly under modern budget and computational constraints (Neale et al. 2017a). For example, analysis of *Pinus taeda* L. (*Pinaceae*) has yielded estimates that upwards of 82% of its 22Gb genome is repetitive (Wegrzyn et al. 2014). It is also thought to be rich in pseudogenes (Kovach et al. 2010).

Such large genome sizes have hampered production of dense SNP datasets across a large number of individuals (Lind et al. 2018). Most recent sequencing efforts in conifers have either used some form of reduced representation sequencing such as restriction-site associated DNA sequencing (i.e., RADseq; reviewed in Andrew et al. 2016 and Parchman et al. 2018), which relies upon relatively few genomic resources, or targeted capture (e.g., Lu et al. 2016; Suren et al. 2016), which requires significant genomic and budgetary resources including the design of capture arrays (but see Puritz and Lotterhos 2018). To capture population-level polymorphism information while minimizing cost, the sequencing of pooled individuals (i.e., pool-seq approaches) has emerged as a cost-effective alternative to sequencing individuals (Gautier et al. 2013; Schötterer et al. 2014). Further, pool-seq can be combined with targeted capture approaches to both reduce cost and sample specific areas of the genome that are *a priori* considered functionally relevant (e.g., Rellstab et al. 2019).

Pool-seq approaches use read counts across pooled individuals to estimate allele frequencies (generally for a single population). A number of studies have empirically evaluated the congruence between individual and pool-seq allele frequency estimates across various taxa (e.g., Futschik & Schlötterer 2010; Rellstab et al. 2013, 2019; Fracassetti et al. 2015). Such studies have led to broad agreement on the accuracy of pool-seq, with best practices following Schlötterer et al. (2014) Of exceptional significance for the estimation of allele frequency from read count data is the proper alignment of reads to the reference. Misalignments, which may be particularly important for exome capture data, can be due to reads from paralogs in the data mapping to the incorrect copy in the reference, or from paralog copies being collapsed into a single sequence in the reference assembly where copies in the data map to this single sequence. These misalignments will skew allele ratios and bias allele frequency estimates downstream. In particular for non-model species with histories of whole genome duplication or gene family expansion, steps must be taken to categorize misalignments so that there are not substantial allele frequency biases in downstream datasets.

Here we harness the multicellular haploid megagametophyte of conifers to aid in mapping and analyzing poolseq data from diploid individuals. We use this pooled exome capture approach for two conifers: coastal Douglas-fir (*Pseudotsuga menziesii* var. *menziesii* (Mirb.) Franco, *Pinaceae*) and jack pine (*Pinus banksiana* Lamb., *Pinaceae*), to evaluate the utility of pool-seq approaches in these systems. We use sequence data from individuals to validate allele frequency estimates of the same individuals in pools, and use haploid sequencing to identify misalignments from paralogous sites to quantify their effects on the congruence between individual and pool-seq estimates (Table 1). Together, these datasets provide a path forward for filtering pool-seq data of this kind, particularly for studies of non-model organisms using a diverged, and potentially fragmented, reference genome. Our methods further highlight a cost-effective means to empirically isolate potentially misaligned paralogs in species with accessible haploid tissue.

**TABLE 1.** Description of datasets used to call SNPs for both Douglas-fir and jack pine. indSeq^†^ and poolSeq datasets for a given species share the same individuals from a single population. The megaSeq dataset consists of haploid megagametophyte tissue from a single individual not included in the indSeq or poolSeq datasets.

| Dataset | Ploidy per sample<br>(number of<br>samples) | SNP<br>Caller | Purpose |
| --- | --- | --- | --- |
| indSeq | 2 (20) | GATK4 | Validate poolSeq allele frequency estimates;<br>calculate read ratio statistics to validate candidate<br>paralog misalignments |
| poolSeq | 2 (20) | VarScan | Compare with indSeq SNP set to<br>determine filtering strategy |
| megaSeq | 1 (1) | VarScan | Identify heterozygous sites as candidates for false<br>SNPs due to misalignment of diverged / duplicated<br>paralogs |

<sup>†</sup> Note that we use camelCase to denote our datasets, and reserve hyphens (e.g., pool-seq) to denote methodologies.

## 2 Methods

### 2.1 Focal Species and Population Sampling

Coastal Douglas-fir (*Pseudotsuga menziesii* var. *menziesii*) is a temperate species occupying primarily coastal habitat along the west coast of North America from Mexico to British Columbia as well as inland habitat in the Cascade and Klamath ranges of Washington, Oregon, and California. It is important to the ecology and economy of many of these forests. Jack pine (*Pinus banksiana*) has the widest distribution of any tree in the vast Canadian boreal forest, stretching from Atlantic Canada into western Alberta and Northwest Territories and is important to the ecology of many of these systems and to the forest industry in some regions.

For both Douglas-fir and jack pine, we sampled 20 individuals for use in individual and pooled sequencing sets from operational reforestation seedlots created from bulk open-pollinated seeds from tens or hundreds of seed parents from a single provenance (see Supplemental Note 1.1). We used a single jack pine seed to extract megagametophyte haploid tissue. For Douglas-fir haploid data, we randomly chose paired-end fastq files from a previously sequenced Douglas-fir megagametophyte taken from a single individual (NCBI SRA accession SAMN0333061, Neale et al. 2017b) to match sequencing effort for jack pine haploid tissue (Supplemental Note 1.2).

### 2.2 Exome Capture Probe Design

The capture probes were designed based on the genes identified using RNA sequencing (RNA-seq) data for Douglas-fir and jack pine. *De novo* transcriptome assembly was performed for each species using RNA-seq reads. For jack pine, RNA-seq reads were sequenced from a sample of needles from a single tree. For Douglas-fir, RNA-seq reads were obtained from two sources: one source was the read sets deposited in NCBI SRA, including SRX1851630 (Little et al. 2016), SRX1286745 (Hess et al. 2016), SRX1341335 (Cronn et al. 2017a), and SRX136240 (Cronn et al. 2017b). The other source was the reads sequenced from two needle samples infected by the fungal pathogen *Phaeocryptopus gaeumannii*, which causes Swiss needle cast disease in Douglas-fir trees.

RNA-seq was conducted by McGill University and Génome Québec Innovation Centre using HiSeq2500 V4 in paired-end 125bp format. The raw reads were processed by the software FASTX-Toolkit (v0.0.13, http://hannonlab.cshl.edu/fastx_toolkit), including clipping the adaptors (-l 25), filtering the artifacts, and keeping the reads with a minimum quality score of 20. The filtered reads were used to perform *de novo* transcriptome assembly using the software Trinity v2.4.0 (--bowtie2, Grabherr et al. 2011). Among the assembled transcripts, only the longest isoforms with a length of at least 300bp for each gene were retained, which were then used as reference to align the reads using the software RSEM (v1.3.0 Li and Dewey 2011). From the expression quantification of transcripts, transcripts with aligned reads and transcript per million 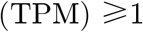 were retained. The completeness of the filtered transcripts was examined using the 1,375 orthologs in the BUSCO (v3.0, Benchmarking Universal Single Copy Orthologs) set of `embryophyta_odb10` (`--evalue 1e-10`, Simão et al. 2015).

To avoid probes spanning exon-intron boundaries, exons were targeted to design probes. Using the software GMAP (v2017-06-20, Wu and Watanabe 2005), the filtered transcripts from Douglas-fir were aligned to the convarietal reference (*Pseudotsuga menziesii var. menziesii* (coastal Douglas-fir; v1.0, Neale et al. 2017b). The jack pine transcripts were aligned to the congeneric loblolly pine (*Pinus taeda*) reference genome (v.1.01, Wegrzyn et al. 2014) as there is no available jack pine reference genome, and both loblolly and jack pine belong to *Pinus* subgenus *Pinus,* the hard pines. Exon sequences with a length of at least 100bp were submitted to Roche NimbleGen for Custom SeqCap EZ probe design.

To evaluate the capture efficiency of the probes, the captured sequences were aligned to reference genomes and numbers of reads on-target, near-target (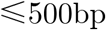 from target regions), and off-target regions were counted using “intersect” function in the software `BEDtools` v2.28.0 (`-f 0.75`, Quinlan and Hall 2010). The depth of captured sequences was counted using “depth” function in the software `SAMtools` v1.3 (`-q 3 0 -Q 20`, Li et al. 2009). The cumulative depth was calculated and plotted using `R` (R Core Team 2018).

### 2.3 DNA Extraction, Library Preparation, and Sequencing

In total, three datasets were created for each of the two species (Table 1) – note that we use camelCase (e.g., poolSeq) to denote our datasets, and reserve hyphens (e.g., pool-seq) to denote methodologies. These datasets included individual sequencing of 20 diploid individuals from a single population (hereafter indSeq), the same individuals pooled together prior to sequencing (hereafter poolSeq), and haploid megagametophyte tissue sequenced from a single individual (hereafter megaSeq). We use the indSeq dataset to validate allele frequency estimates from our poolSeq data, and the megaSeq data to probe our data for heterozygote SNPs (i.e., potential false-positive SNPs) caused by the misalignment of diverged paralogs that could affect our allele frequency estimates (Table 1; see also *§2.6 / Comparison, of Sequencing Approaches).*

For each dataset we extracted DNA from either diploid needle tissue or haploid megagametophyte tissue (see Supplemental Note 1.3). From these extractions, approximately 100ng of DNA from each individual or pooled DNA sample was used for a barcoded (Kapa, Dual-Indexed Adapter Kit) library with an approximately 350-bp mean insert size. SeqCap library preparation was performed using custom Nimblegen SeqCap probes (described above in *§2.1*) according to the NimbleGen SeqCap EZ HyperCap Workflow User’s Guide Ver 2 (Roche Sequencing Solutions, Inc., CA USA). Following capture, each library was sequenced in a 150bp paired-end format on an Illumina HiSeq4000 instrument at the Genome Quebec Innovation Centre at McGill University, Montreal, Canada.

### 2.4 Bioinformatic SNP Calling Pipelines

Raw paired-end sequence reads from all datasets were trimmed with `fastp` (v0.19.5, Chen et al. 2018) by trimming reads that did not pass quality filters of less than twenty N’s, a minimum mean Phred quality score of 30 for sliding windows of five base pairs, and a final length of 75 base pairs with no more than 20 base pairs called as N (`-n 2 0 -M 3 0 -W 5 -l 75 -g -3`). Trimmed reads were mapped with `bwa-mem` (v0.7.17, Li & Durbin 2009) to reference assemblies; we mapped jack pine to the loblolly reference (v2.01, Wegrzyn et al. 2014) and Douglas-fir to the convarietal reference (v1.0, Neale at al. 2017b). The resulting `.sam` files were converted to binary with `SAMtools` v1.9 (`view`, `sort`, `index`; Li et al. 2009) and subsequently filtered for proper pairs and a mapping quality score of 20 or greater (`view -q 20 -f 0x0002 -F 0x0004`). Using `Picardtools` v2.18.9 (http://picard.sourceforge.net), read groups were added and duplicates subsequently removed from filtered bam files.

We then called SNPs using the Genome Analysis Toolkit (`GATK` v4.1.0.0; McKenna et al. 2010) for indSeq data, and `VarScan` (v2.4.3; Koboldt et al. 2012) for both poolSeq and megaSeq datasets (Table 1) for comparisons since future datasets that stem from a larger project will all be poolSeq (and we will therefor only be using `VarScan`). For SNPs called with `GATK4`, we used `HaplotypeCaller` (`--genotyping-mode DISCOVERY -ERC GVCF`) and `GenotypeGVCFs`. We then filtered data with `SelectVariants` (`--select-type-to-include SNP`), `VariantFiltration` (`--filter-expression “QD < 2.0 || FS > 60.0 || MQ < 40 || MQRankSum < -12.5”`), and finally with `vcftools` v0.1.14 (`-- maf 0.00 –minGQ 20 –max-missing 0.75`; Danecek et al. 2011). BQSR was not carried out in our analysis due to the lack of a high-quality reference set of SNPs for our species. Note that no further filtering (e.g., for depth) was done for this initial baseline filtering strategy (further filtering is described in *§3.4*).

Before calling SNPs with VarScan, we first realigned indels with `GATK` 3.8 (McKenna et al. 2010) – `RealignerTargetCreator` then `IndelRealigner` – and then passed a `SAMtools mpileup` object directly to `VarScan::mpileup2cns` with a minimum coverage set to 8, *p*-value < 0.05, minimum variant frequency of 0.00, ignoring variants with >90% support on one strand, a minimum average genotype quality of 20, and a minimum allele frequency of 0.80 to call a site homozygous (`--min-coverage 8 --p-value 0.05 --min- var-freq 0.00 --strand-filter 1 --min-avg-qual 20 --min-freq-for-hom 0.80`). Output was then filtered with a custom python (v3.7, www.python.org) script to filter out indels, keep only biallelic loci, and to ensure a genotype quality score > 20. From the megaSeq data, we then isolated heterozygous SNP calls (hereafter megaSNPs) that represent errors in genotype calling given the haploid nature of the tissue sequenced – to keep only heterozygous calls, we ignored any biallelic cases where only the non-reference allele was called. Such apparent SNPs are likely false due to misalignments. We have published our complete SNP calling pipelines in publicly available repositories (https://GitHub.com/CoAdapTree/gatk_pipeline and ./varscan_pipeline).

### 2.5 Validation of megaSNPs as indicators of paralogy artifacts

To check whether heterozygous sites (megaSNPs) called from `VarScan` megaSeq are following expectations of patterns from paralogs, we investigated read ratio deviations from a binomial expectation for these `VarScan` megaSNP sites at the same sites in our `GATK` indSeq data using heterozygous diploid individuals (*sensu* McKinney et al 2017; see also Rellstab et al. 2019). For true positive SNPs, heterozygous diploid individuals should have, on average, an even ratio of reference (REF) and alternative (ALT) read counts. If the SNP is due to a bioinformatic error arising from the misalignment of paralogs (i.e., a false positive SNP), the read ratio will differ significantly from this expectation when there is a SNP at a given position in only one paralog copy (McKinney et al. 2017). Similarly, if there is a fixed difference at a given position between two copies, then all individuals in a population will present as heterozygotes with balanced read counts for REF and ALT at that site. If we are sequencing (and then post-hoc correctly identifying) paralogs in our poolSeq data using megaSNP sites, misalignment of either duplicated or diverged paralogs will cause read ratio deviations in these loci (and affect allele frequency estimates from poolSeq, and downstream analyses), which we should be able to detect in our indSeq data. As described by McKinney et al. (2017), subsequent to whole-genome duplication during the rediploidization phase as homeologous chromosomes diverge, tetrasomically inherited sets of paralogs (duplicates) organize into distinct disomic loci (diverged duplicates).

We calculated these read ratio statistics for sites within the intersection of (1) megaSNPs, indSeq, and poolSeq SNPs, and (2) poolSeq and indSeq SNPs alone; hereafter intersections I1 and I2. The purpose of (1) is to see how paralogs could affect our poolSeq data (leveraging information in our indSeq data to do so), and of (2) is to visualize the potential influence of paralogs in our data independent of sites identified as megaSNP sites, as well as to compare poolSeq allele frequency estimates with those estimated from the indSeq dataset. For these sites, we queried the indSeq data to record the frequency of heterozygous individuals (*H*), the allele depth ratio 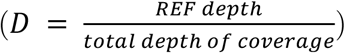, and the deviation of allele depth from expectation (REF depth – 0.5*total depth) standardized by properties of a binomial distribution with *n* = depth of coverage, and *p* = 0.5 (i.e., the *z-* score for the allele ratio deviation) following McKinney et al. (2016, 2017). We compare our results with simulations carried out by McKinney et al. 2017. We used custom `python` code to replicate the methods of McKinney et al. (2016), which is available online at https://github.com/CoAdapTree/testdata_validation/003_testdata_validate_megaSNPs.ipynb.

### 2.6 Comparison of sequencing approaches

To study the utility of our pooled exome capture approach, we compared estimates of allele frequency from our indSeq data with estimates from our poolSeq data. To do so, we took the baseline-filtered SNPs from poolSeq and indSeq (see *§2.4 | Bioinformatic SNP Calling Pipelines*) and identified common SNPs (i.e., intersection I2). To quantify and visualize congruence between allele frequencies estimated with these methods, we report Pearson’s correlation coefficient, plot histograms to visualize the congruence across the minor allele frequency (MAF) spectrum, and further plot 2D histograms to visualize congruence of allele frequency estimates. To visualize how filtering poolSeq SNPs affects the congruence between indSeq and poolSeq allele frequency estimates, we plot the allele frequency differences between methods (hereafter *AFdiff,* calculated as the difference in allele frequency methods of poolSeq and indSeq: *poolSeq_AF_* – *indSeq_AF_*) against poolSeq MAF, poolSeq depth of coverage, *H*, and the **z**-score of read ratio deviation.

## 3 Results

### 3.1 Sequencing, Mapping, and Probe efficiency

Sequencing of the prepared libraries resulted in high quality datasets, with the average base quality above 30 before trimming having a mean of 86.99% across datasets and species, and a mean of 89.43% after trimming (Table S1). The number of sequenced reads varied across datasets but was similar within datasets (on average 405 million reads for indSeq, 130 million reads for poolSeq, 202 million reads for megaSeq). Mapping rates generally reflected the phylogenetic relationship between the sequenced individuals and the reference used, where rates were high for all coastal Douglas-fir datasets mapping to the convarietal reference (mean 85.11%) and relatively lower rates for jack pine datasets mapping to the congeneric *Pinus taeda* reference (mean 35.36%; Table S1).

The BUSCO analysis to assess completeness of transcripts used in exome-capture probe design resulted in recovery of 87% of the 1,375 BUSCOs in Douglas-fir transcripts, including 70% complete and single-copy BUSCOs, 2% complete and duplicated BUSCOs, and 15% fragmented BUSCOs. For jack pine transcripts, 93% of the BUSCOs were recovered, including 85% complete and single-copy BUSCOs, 2% complete and duplicated BUSCOs, and 6% fragmented BUSCOs. We aligned the transcripts to reference genomes to select exons and design probes. The final capture probe size are 41 Mbp for jack pine (design name: 180215_jackpine_v1_EZ_HX1) and 39 Mbp for Douglas-fir (design name: 80215_DOUGFIR_V1_EZ), corresponding to 32,208 genes in jack pine and 37,787 genes in Douglas-fir.

We counted the number of captured reads on-target, near-target (≤500bp from target), and off-target regions for indSeq and poolSeq samples. The poolSeq samples had more reads than the indSeq samples and off-target regions had the most aligned reads (Figure 1). Reliable SNP calling is dependent on sequencing depth, so we calculated the cumulative numbers of bases on different regions. For Douglas-fir poolSeq, we obtained over 40 million bases in target regions with at least 20X sequencing depth (Figure 2A). For jack pine poolSeq, we obtained over 30 million bases in on-target regions with at least 20X sequencing depth (Figure 2B). Sequencing depths in near- and off-target regions were dramatically diminished compared to the on-target regions.

**FIGURE 1.**
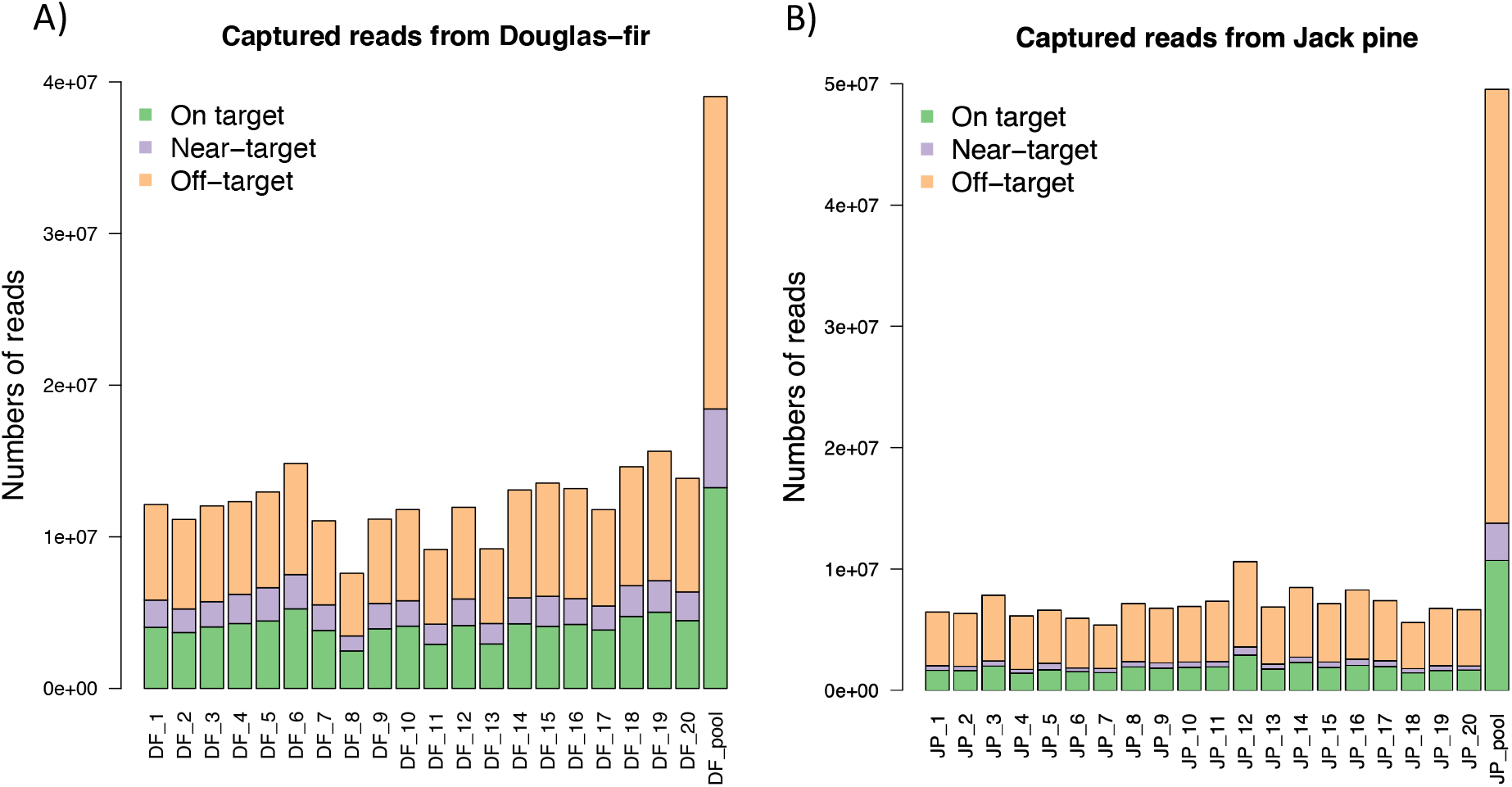
Numbers of captured reads from Douglas-fir (A) and jack pine (B) that mapped on target, near-target (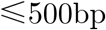 from target) and off-target regions. On the x-axis, from left to right, the first 20 bars represent indSeq samples, and the last bars represents the poolSeq samples.

**FIGURE 2.**
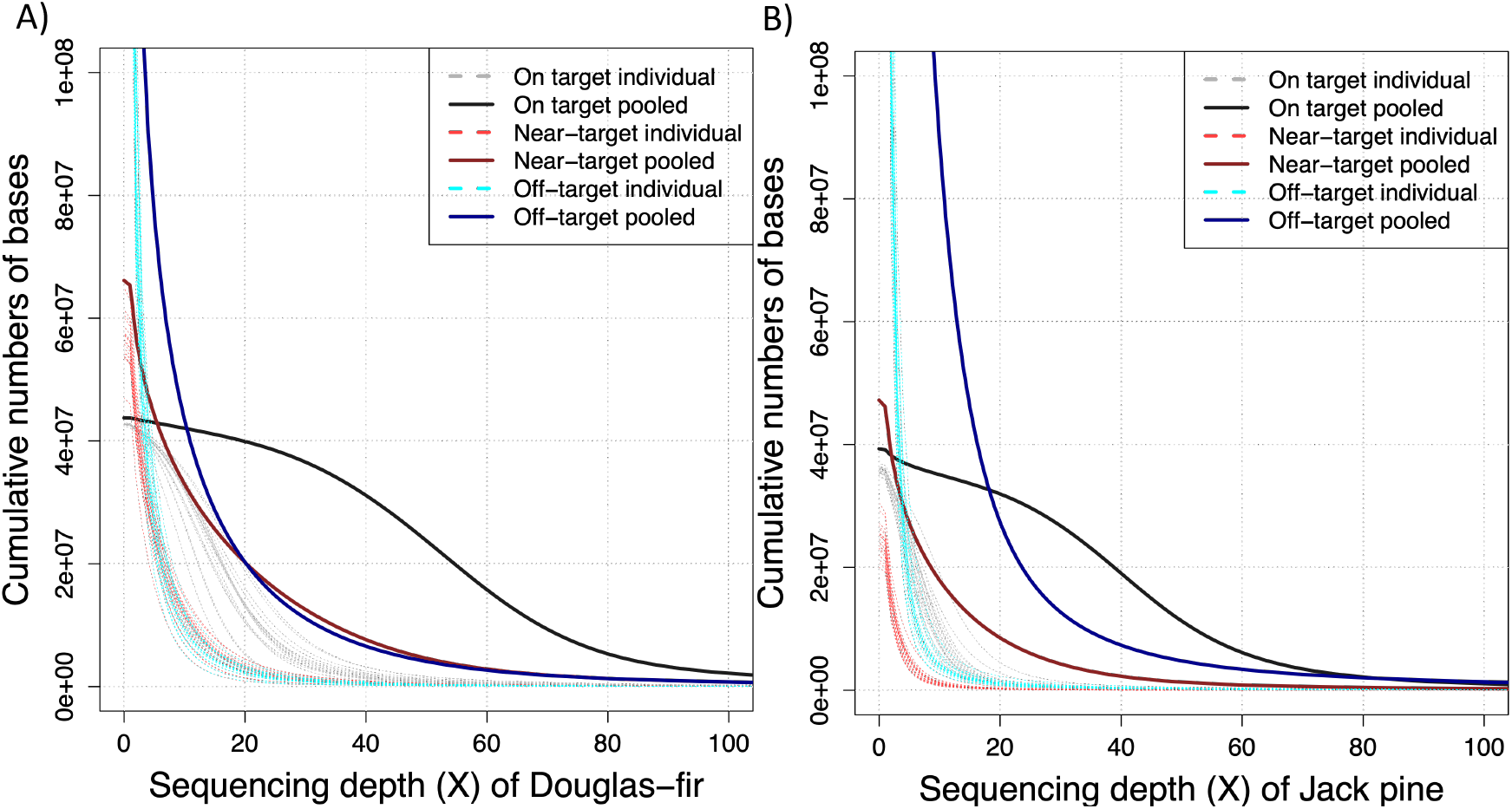
Cumulative numbers of bases in Douglas-fir (A) and jack pine (B) on target, near-target (≤500bp from target), and off-target regions. Dashed lines represent the cumulative numbers of bases in each of 20 indSeq samples. Solid lines represent the cumulative numbers of bases in the poolseq sample.

### 3.2 SNP calling

The total number of SNPs after baseline filtering varied across datasets and species (Table 2). Douglas-fir generally had a higher number of SNPs called than jack pine, except for poolSeq data. However, there were more jack pine megaSNPs intersecting with poolSeq (25,500 SNPs) and indSeq (7,408 SNPs) than for Douglas-fir datasets (respectively 825 SNPs and 293 SNPs). Given that megaSNPs are cases where a heterozygote call was made from a haploid sample and are therefore indicators of bioinformatic-paralogy errors, this suggests that the error rate is much higher in jack pine. In total, several hundred thousand SNPs were found in the intersection of poolSeq and indSeq for Douglas-fir (636,279 SNPs) and jack pine (255,706 SNPs; Table 2).

**TABLE 2.** Output of SNPs from the conifer datasets. The intersection across all three baseline-filtered datasets were 7,006 SNPs for jack pine (JP) and 248 SNPs for Douglas-fir (DF).

| Dataset | Species | Baseline-filtered SNPs | Baseline Intersecting SNPs |  |
| --- | --- | --- | --- | --- |
|  |  |  | poolSeq | megaSeq<sup>†</sup> |
| indSeq | DF | 1,526,554 | 636,279 | 293 |
|  | JP | 377,080 | 255,706 | 7,408 |
| poolSeq | DF | 1,601,285 | - | - |
|  | JP | 3,686,528 | - | - |
| megaSeq<sup>†</sup> | DF | 398,774 | 825 | - |
|  | JP | 32,751 | 25,500 | - |

<sup>†</sup> These numbers reflect only heterozygous SNPs (i.e., megaSNPs)

### 3.3 Validation of megaSNPs as indicators of paralogy artifacts

Upon inspection of our intersecting sets, patterns expected for duplicated and diverged duplicate paralogs (McKinney et al. 2017) were apparent in both intersection I1 (megaSNPs, indSeq and poolSeq SNPs) and intersection I2 (indSeq and poolSeq SNPs), but megaSNPs did not generally typify patterns expected from non-duplicated (singleton) genes. For instance, SNPs in duplicated genes should be most distinct from SNPs in singletons when the derived allele is at intermediate frequency, and diverged duplicates most distinct from singletons when the derived allele is fixed (Figure 3A). Sites consistent with expectations for singleton, duplicates, and diverged duplicates were apparent from intersection of poolSeq and indSeq sites (i.e., intersection I2; Figure 3D-E), while the indSeq sites intersecting with candidate paralog sites (megaSNPs, i.e., intersection I1) displayed elevated levels of heterozygosity as expected from paralogs (Figure 3B-C). Indeed, patterns of deviated allele ratios were also seen in our data (Figures S1 D-E, S2 D-E), where the vast majority of megaSNP sites were considerably different than the 1:1 read ratio expected of heterozygous diploids (Figures S1B-C, S2B-C) as would otherwise be expected for singletons (Figures S1A, S2A). Lastly, when considering the standardized allele ratio deviation (*z*-score) we recover the same patterns of point clouds classified by McKinney et al. (2017). We observe a dense set of SNPs around the *z*-score of 0.0 for *H* values of 0.0-0.6 (Figure 4D-E) expected from singleton sites (Figure 4A), another set of SNPs with elevated *H* and/or *z*-score (Figure 4B-C) that is expected from duplicate loci (Figure 4A), and a third set of SNPs with *H* > 0.9 (Figure 4B-C) that is expected for diverged duplicates (compare to Figures 5 and 8 in McKinney et al. 2017).

**FIGURE 3.**
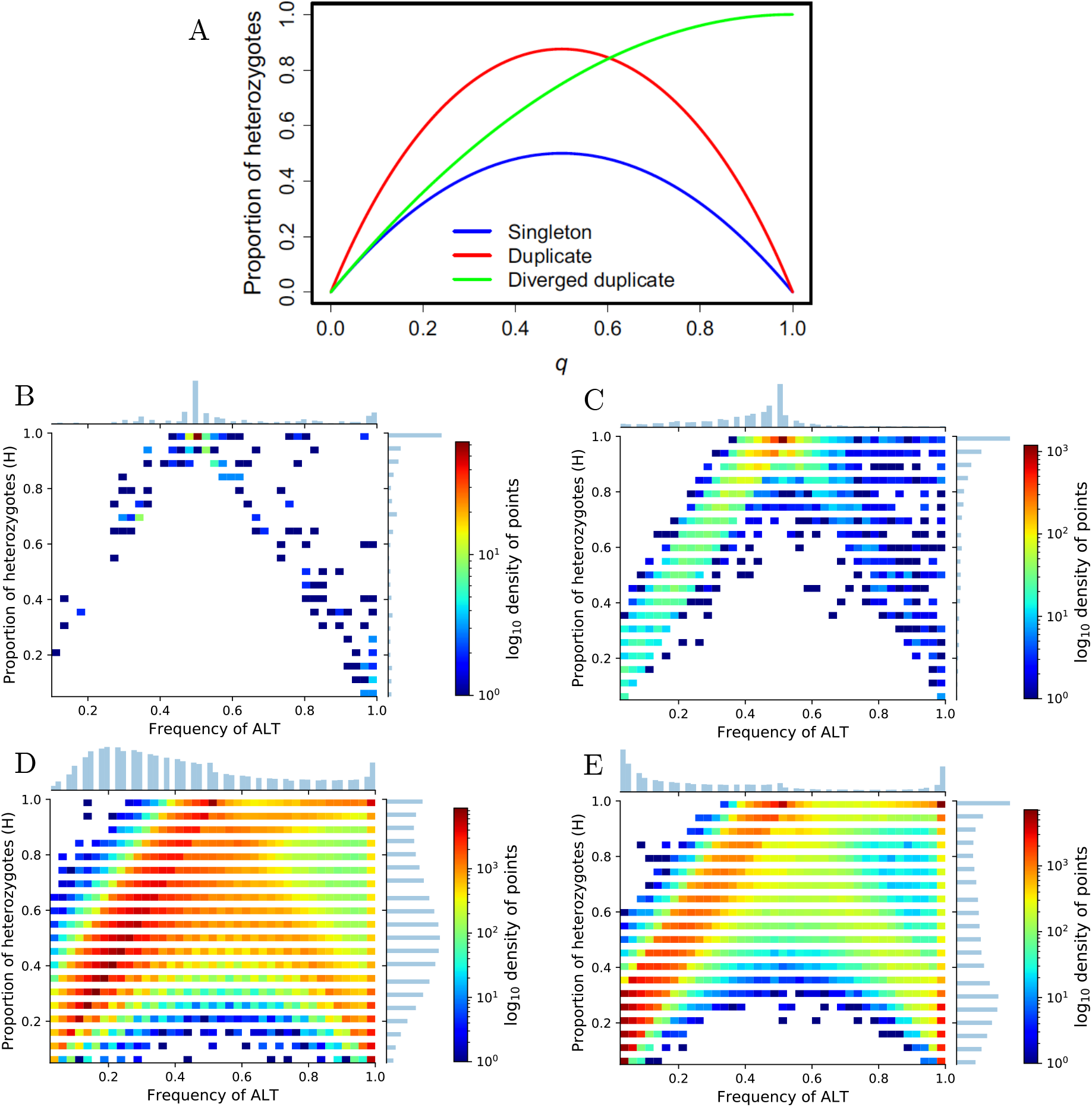
The proportion of heterozygotes, *H* and the alternative (ALT) frequency calculated from indSeq data distinguish paralog misalignments according to expectations (A, Fig. 4 from McKinney et al. 2017), and empirically for Douglas-fir (B, D) and jack pine (C, E). B-C) Empirical distribution of megaSNP sites (candidate paralog sites identified as heterozygote calls from haploid tissue) calculated using indSeq data for those sites that were also called in poolSeq data (i.e., intersection I1). D-E) Empirical distribution of intersection I2 (indSeq and poolSeq intersection) calculated using indSeq data. Note color scale changes for each figure to accentuate patterns in the data. Frequency of ALT was binned for visualization purposes. Code to create these figures is available online: https://github.com/CoAdapTree/testdata_validation

**FIGURE 4.**
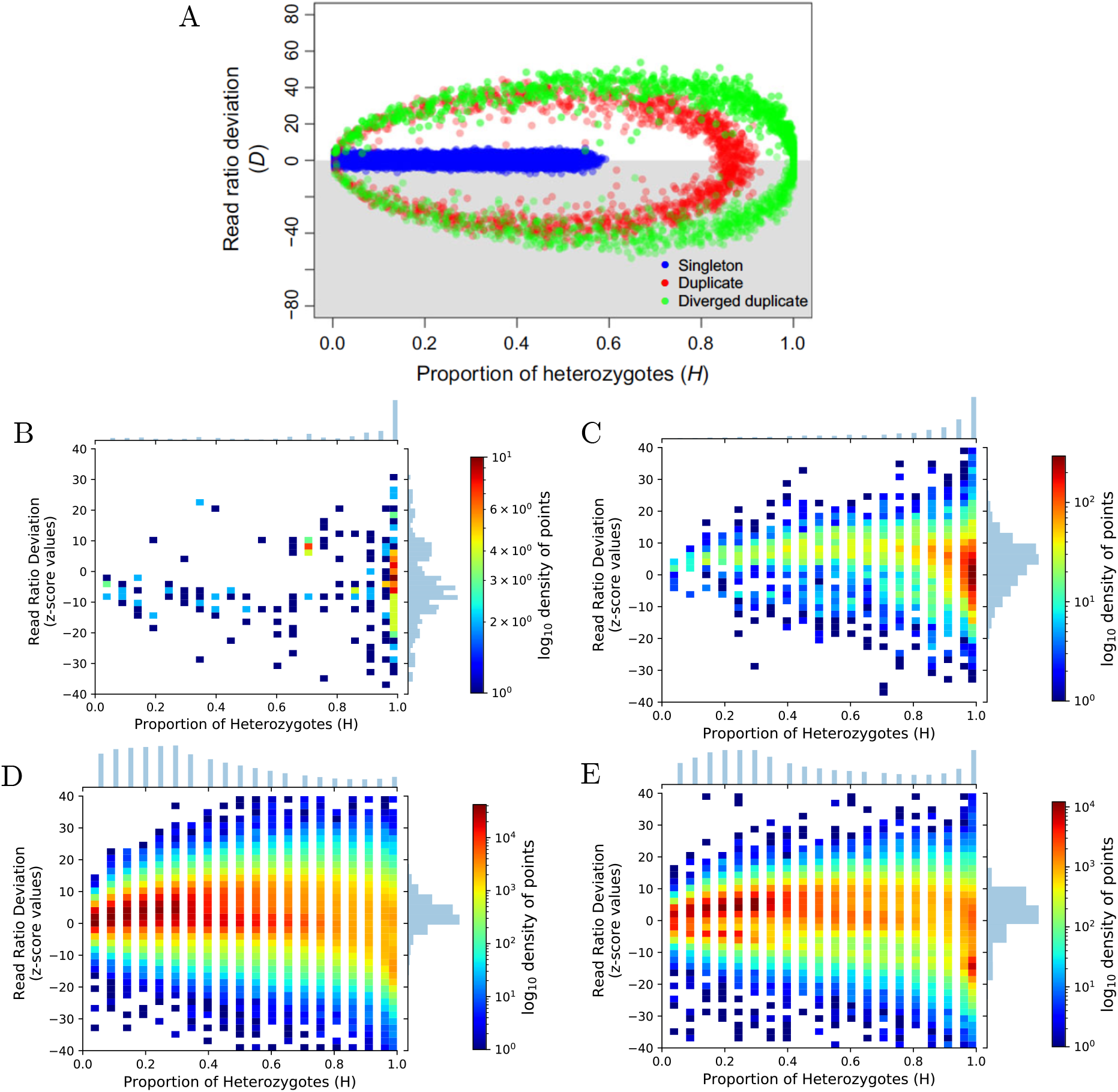
Standard deviation of read ratio (z-score) and the percentage of heterozygotes (*H*) calculated from indSeq data distinguish paralog misalignments according to expectations (A, Fig. 4 from McKinney et al. 2017), and empirically for Douglas-fir (B, D) and jack pine (C, E). B-C) Empirical distribution of megaSNP sites (candidate paralog sites identified as heterozygote calls from haploid tissue) calculated using indSeq data for those sites that were also called in poolSeq data (i.e, intersection I1). D-E) Empirical distribution of intersection I2 (indSeq and poolSeq intersection) calculated using indSeq data. As with 3A, the distribution found in the gray box in 4A will be found in the upper white panel because we used the reference allele instead of randomly choosing the allele. Note color scale changes for each figure to accentuate patterns in the data. Code to create these figures is available online: https://github.com/CoAdapTree/testdata_validation

**FIGURE 5.**
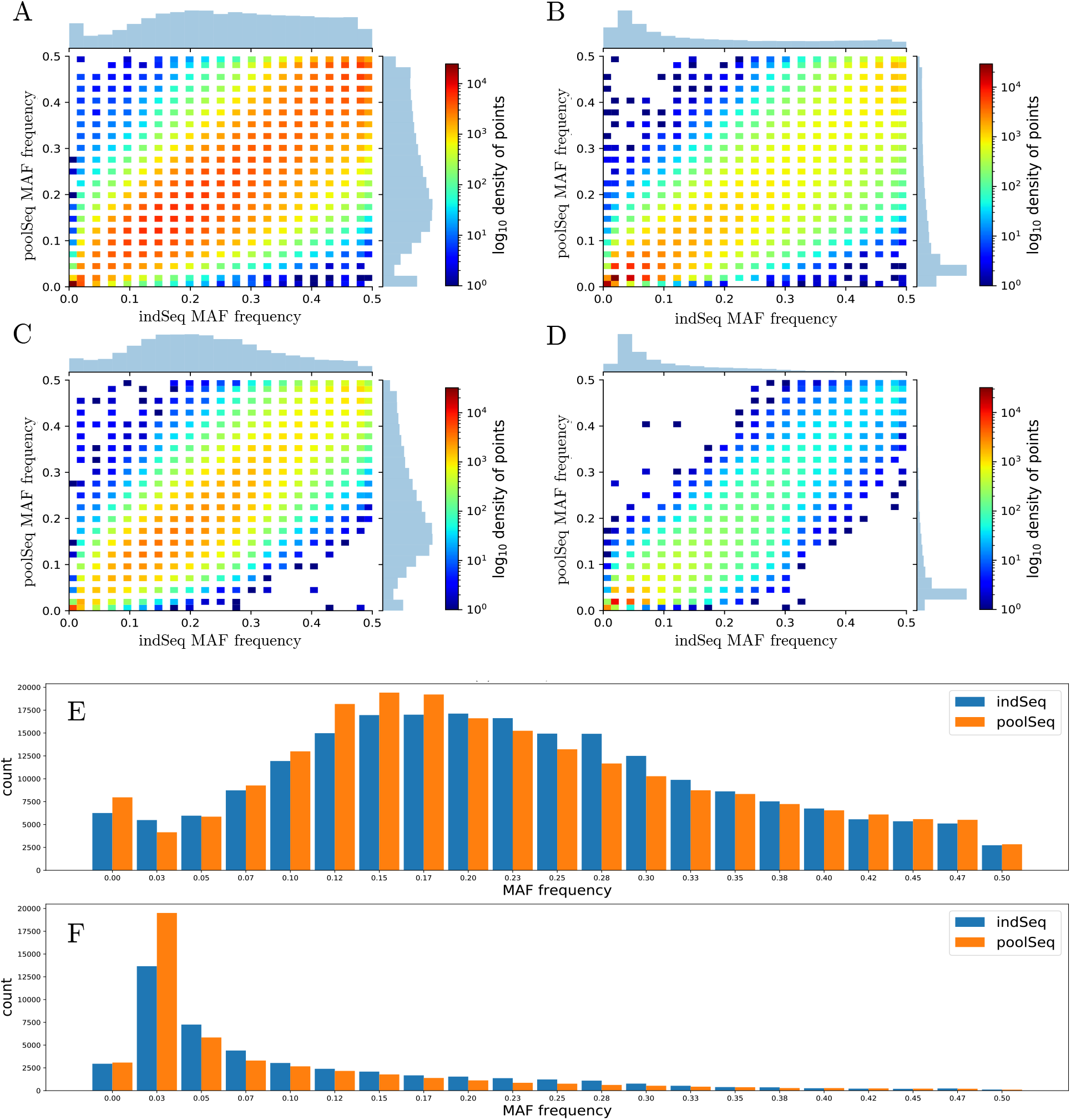
Congruence between indSeq and poolSeq (respectively x- and y-axes, A-D) minor allele frequency (MAF) estimates from Douglas-fir (A, C, E) and jack pine (B, D, F). A-B) 2D histogram of baseline-filtered intersection between indSeq and poolSeq (i.e., intersection I2). C-D) 2D histogram for SNPs after filtering intersection I2 for megaSNP sites, *H* > 0.6, *abs*(z-score) > 10, and indSeq sites with > 20% missing data. E-F) Congruence of minor allele frequency spectra from SNPs in (C) and (D). Color scale is standardized to visualize differences in density between filtering steps. Minor allele frequency was binned for visualization purposes. Code to create these figures is available online: https://github.com/CoAdapTree/testdata_validation

### 3.4 Comparisons of sequencing approaches

Loci within the intersection of baseline-filtered indSeq and poolSeq datasets (i.e., intersection I2) showed a strong positive association between allele frequencies estimated from indSeq and poolSeq (Pearson’s *r* = 0.9760, *p* = 0.0000 for jack pine; Pearson’s *r* = 0.9483, *p* = 0.0000 for Douglas-fir; Figure 5A-B). Comparison of the MAF spectrum from these estimates also revealed good agreement by frequency bins (Supplemental Figure S3). After exploring various filtering strategies (see Supplemental Note 1.4), we applied filters that showed a positive effect on congruence between allele frequency estimates in our data (removing megaSNP sites and indSeq sites with *H >* 0.6); that resulted in removing sites with extreme values of *AFdiff* (removing indSeq sites with *z*-score > 10; filtering *H* > 0.6 alone also had this effect); and, finally, that gave us the best estimate of indSeq allele frequency – our standard of comparison – and thus the best impression of the performance of our poolSeq approach (removing indSeq sites with > 20% missing data). The correlation of allele frequencies estimated from indSeq and poolSeq data increased after this filtering (Pearson’s *r* = 0.9907, *p* = 0.0000 for jack pine; Pearson’s *r* = 0.9732, *p* = 0.0000 for Douglas-fir) with relatively fewer sites with extreme differences in the estimates from each method (see top-left and bottom-right corners of 2D histograms; Figure 5A-D). While some differences remain in the estimates of the minor allele frequency spectrum (Figure 5E-F), these two methods largely agree, suggesting a robust poolSeq dataset for further biological hypothesis testing.

## 4 Discussion

Validation of pool-seq approaches commonly involves model organisms with complete or near-complete chromosome-scale reference genomes (e.g., see Table 1 in Rellstab et al. 2013). However, there are few studies that explore this congruence in non-model organisms such as conifers with large and highly fragmented reference genomes, and histories of whole genome duplications and gene family evolution. Here we show that combining exome capture and pool-seq can be an efficient method for quantifying genetic polymorphisms in two such species, and that heterozygous SNPs from haploid data (megaSNPs) consistently uncover sites with patterns expected from the misalignment of paralogs (Figures 3-4, S1-S2). Further, we appear to uncover more false-positive variation in jack pine than in Douglas-fir (Table 2). Concordance of allele frequency estimates from baseline-filtered indSeq and poolSeq datasets (i.e., intersection I2) was strong (*r* > 0.948), and increased with increased filtering, including the filtering of potential false-positive sites (*r* > 0.973, Figure 5). These values are well within the range expected from previous pool-seq studies (Table 1 in Rellstab et al. 2013), and in some cases perform better than these model organisms.

Despite their role in adaptation and speciation (Lynch & Conery 2000; Allendorf et al. 2015), the exclusion of potentially paralogous sites from next generation sequencing datasets is commonplace due to the difficulty in differentiating genetic polymorphisms from differences present among copies from single or diverged gene families (Dou et al. 2012; Hohenlohe et al. 2012; Dufresne et al. 2014). There are several methods by which to detect such problematic sites, such as filtering by coverage (Dou et al. 2012), disomic models such as Hardy-Weinberg proportions (Hohenlohe et al. 2011; Catchen et al. 2013; Chen et al. 2014), or gene annotation, though there are several shortcomings (see descriptions of these shortcomings in Table 1 of McKinney et al. 2017). When individual sequencing data is available for the same individuals or populations, such information can be used to isolate potentially paralogous sites from pool-seq exome capture studies (e.g., Rellstab et al. 2019). However, a potentially cost-saving alternative would be to sequence the haploid tissue of a single individual (if available). Even so, there may be reduced power to detect recently diverged paralogs (i.e., when derived alleles are at low frequency and therefore not readily detected in a single individual). As such, heterozygous SNPs called from our haploid data (megaSNPs) allowed us to identify variation from paralogous misalignments that infrequently displayed patterns expected of singleton gene copies. Indeed, high quality heterozygous calls from haploid sequencing are a reliable method for identifying misalignments due the known monoallelic state of the sequenced site (Limborg et al. 2016). While metrics from sequences of individuals are reliable (McKinney et al. 2017), they can falsely flag potentially paralogous sites as SNPs due to the stochastic nature of the sequencing process and may result in the exclusion of biologically meaningful information.

As pointed out by Rellstab et al. (2019), use of exome capture in many *Pinaceae* species will require particular care to exclude potentially paralogous sites from downstream analysis to avoid biased results. This is particularly true for pool-seq datasets relying on read counts for allele frequency estimation or population genetic inferences such as genotype-environment association (e.g., as implemented in baypass, Gautier et al. 2015). While individually sequenced datasets may be one path forward to identifying such problematic sites (as in Rellstab et al. 2019, Shu & Moran 2020), it will be cost-prohibitive for large projects with many populations. As such, the sequencing of sufficient quantities of DNA from haploid gametophyte tissue available for some plants, including conifers, seedless vascular plants, and bryophytes, offers an alternate path forward to balance sequencing cost and data reliability.

## Acknowledgments

The CoAdapTree project is funded by Genome Canada (241REF), Genome BC and 16 other sponsors (http://coadaptree.forestry.ubc.ca/sponsors/), including Genome Alberta, Génome Québec, the BC Ministry of Forests, Natural Resources Operations and Rural Development, and Compute Canada. We thank Centre d’expertise et de services Génome Québec for sequencing service, University of Calgary Information Technologies for system support, and Dr. Pia Smets and Christine Chourmouzis for technical assistance. We also thank CoAdapTree Scientific Advisory Board members John Davis, Matias Kirst, and Graham Coop for their guidance. Dr. Sam Yeaman is also funded by the Natural Sciences and Engineering Research Council of Canada (NSERC) and Alberta Innovates. Dr. Sally Aitken is funded by an NSERC Discovery Grant. The funding bodies did not have any role in the design of the study, collection, analysis, or interpretation of data in writing the manuscript.

## Data Accessibility

At the time of submission, all code needed to process raw sequence data through figure generation has been made available on GitHub^1^,^2^. Commands used in probe design used default arguments unless otherwise specified. Upon acceptance, a ‘release’ of all code included with submission will be created and remain on GitHub, and will be subsequently archived on DataDryad.org.

Genomic and RNA sequence data will be deposited on the Sequence Read Archive of the National Center for Biotechnology Information (NCBI SRA) as well as the Gene Expression Omnibus. Nimblegen SeqCap probes are available for jack pine under design name 180215_jackpine_v1_EZ_HX1, and for Douglas-fir under design name 80215_DOUGFIR_V1_EZ.

## Author Contributions

SA and SY obtained funding for the study. SA, SY, BL, and ML conceived the research design with contributions from TB. DOV generated the sequence data which BL analyzed, with contributions from ML, PS, and TB. ML designed probes with contributions from SY. BL and ML designed SNP calling pipelines, which were coded by BL and reviewed by ML. BL wrote the manuscript with contributions from ML. All authors contributed to the editing of this manuscript.

## Footnotes

1 https://github.com/CoAdapTree/varscan_pipeline

2 https://github.com/CoAdapTree/testdata_validation

