## Supplemental Information for "Haploid, diploid, and pooled exome capture recapitulate features of biology and paralogy in two non-model tree species"

October 6, 2020

^1^Centre for Forest Conservation Genetics and Department of Forest and Conservation Sciences, University of British Columbia, Vancouver, BC, Canada

^2^Department of Biological Sciences, University of Calgary, Calgary, AB, Canada

^3^Biodiversity Research Centre, University of British Columbia, Vancouver, BC, Canada

*corresponding author

^#^equal contribution

**Running Title**: *Validating pool-seq data in conifers*

**Keywords**: pool-seq, non-model, paralogy, *Pinaceae*, exome-capture

**Corresponding Author**

Brandon Lind

2424 Main Mall

3027 Forest Science Centre

University of British Columbia

Vancouver, BC, Canada

### 1 | Supplemental Text

#### 1.1 Douglas-fir and jack pine seed lot sourcing

Douglas fir seeds originating in California (seed lot ID 02745, US seed zone 525 – California; latitude 39.38, longitude -120.67, elevation 914m) were obtained from the USDA Forest Service Placerville Nursery, Camino, California, USA, while jack pine seeds originating in Saskatchewan, Canada (NTSC number 950112; Saskatchewan, Canada, latitude 54.0833, longitude -107.25, elevation 492m), were received from National Tree Seed Centre (NTSC), Canadian Forest Service, Fredericton, New Brunswick, Canada.

Jack pine seed used as a source of megagametophyte tissue was obtained from NTSC (NTSC number 20002284.0).

#### 1.2 Douglas-fir haploid sequencing data

We used previously sequenced Douglas-fir haploid megagametophyte tissue from a single individual (NCBI SRA accession SAMN03333061; Neale et al. 2017c) by randomly choosing paired-end fastq files from the accession to match the sequencing effort put forth towards jack pine haploid tissue. In doing so we used the following files in our Douglas-fir megaSeq SNP calling pipeline:

1. SRR2027119_1.fastq.gz and SRR2027119_2.fastq.gz
2. SRR2027122_1.fastq.gz and SRR2027122_2.fastq.gz
3. SRR2027123_1.fastq.gz and SRR2027123_2.fastq.gz
4. SRR2027124_1.fastq.gz and SRR2027124_2.fastq.gz
5. SRR2027125_1.fastq.gz and SRR2027125_2.fastq.gz
6. SRR2027128_1.fastq.gz and SRR2027128_2.fastq.gz

#### 1.3 DNA extraction protocols

*DNA extraction megagametophyte tissue*

For Douglas-fir, please see Supplemental Note 1.2.

For jack pine, seed was incubated for one hour in a Petri dish on a wet filter paper at room temperature to hydrate the seed tissues. The next step was to freeze the seed for a couple of hours at -20°C and then let it defrost for 30 minutes at room temperature. To avoid the dirt and resin remains on the seed coat to contaminate the inner tissues, the seed coat was scraped with a surgical blade and all dirt was removed by washing the seed in sterile water.

The outer seed coat was removed using a surgical blade under a binocular stereo microscope. The remaining seed content was transferred to a new microscope slide and washed in a drop of 100% ethanol in order to detach the megagametophyte tissue from the inner seed coat. Precise removal of the inner coat is an essential step because this fine membrane-like coat is a diploid tissue and therefore, there is a high risk of cross-contamination between the tissues. The megagametophyte was cut lengthways to remove the embryo.

Megagametophyte tissue (approx. 3mg) was placed in a 1.5 ml tube and kept at -20°C until further proceeding with the DNA extraction. We used the NucleoSpin Plant II Mini kit (Macherey–Nagel GmbH & Co. KG, Germany) to extract DNA, following modifications recommended by García and Escribano-Ávila (2016).

*DNA extraction from needle tissue*

Seeds were stratified at 4°C for 4 weeks, and subsequently germinated and grown in a Conviron growth chamber with a simulated climate corresponding to the mean annual temperature of 6 °C. Young needle tissue was collected from 20 Douglas-fir and 20 jack pine seedlings after 11 weeks of growth. DNA was extracted using the Nucleospin 96 Plant II Core kit (Macherey–Nagel GmbH & Co. KG, Germany), automated on an Eppendorf EpMotion 5075TM liquid-handling platform. Twenty individual DNA samples from each species were normalized at 10ng/μl and a pooled sample was prepared by combining equimolar amounts of 20 individual DNA samples prior to library preparation.

#### 1.4 SNP filtering

The difference between poolSeq and indSeq estimates ($AFdiff$) generally did not follow any pattern with the poolSeq estimated MAF, except for the lowest MAF bin (Supplemental Figure S4), suggesting that accuracy of poolSeq estimates in our data (assuming indSeq to be accurate) is generally not strongly affected by the frequency of the minor allele. Indeed, removing poolSeq SNPs from intersection I2 with MAF < 0.05 decreased the correlation between datasets (Pearson’s $r$ = 0.9373, $p$ = 0.0 for jack pine; Pearson’s $r$ = 0.9230, $p$ = 0.0 for Douglas-fir).

Removing megaSNP sites from intersection I2 increased the relationship between indSeq and poolSeq allele frequency estimates, though not substantially (Pearson’s $r$ = 0.9776, $p$ = 0.0 for jack pine; Pearson’s $r$ = 0.9483, $p$ = 0.0 for Douglas-fir) and not to the point that could be seen visually with 2D histograms (Supplemental Figures S6A-B). Similarly, removing from intersection I2 any poolSeq SNP with a depth of coverage less than 30 also increased the correlation between indSeq and poolSeq frequency estimates, but not substantially. From this point forward, we consider filtering after removing megaSNP sites from the I2 intersection (hereafter I2B).

The read ratio $z$-scores estimated from indSeq data in I2B revealed that as the $z$-score becomes more extreme, the absolute differences in allele frequency between datasets ($AFdiff$) generally increase (Supplemental Figure S5). However, filtering I2B to remove any site with a $z$-score > 10 decreased the correlation between datasets but not substantially (Pearson’s $r$ = 0.9759, $p$ = 0.0 for jack pine; Pearson’s $r$ = 0.9449, $p$ = 0.0 for Douglas-fir); we also observed similar patterns for increasing thresholds for $z$-score. Even so, we were able to observe via 2D histograms that sites with the most extreme differences in estimated MAF dropped out (Supplemental Figures S6C-D).

When plotted against $H$ (the percentage of heterozygotes found in indSeq data), the difference in allele frequency estimates, $AFdiff$, generally had larger variation at high levels of heterozygote individuals, and less so with a lower value of $H$ (Supplemental Figure S7). As a result, removing sites with a value of $H > 0.6$ (as in e.g., Rellstab et al. 2019) increased the correlation of allele frequency estimates between datasets, and we again observed that sites with the highest discordance in allele frequency estimates began to drop out of our data (Supplemental Figure S6E-F).

### 2 | Supplemental Tables and Figures.

| **Table S1**. Trimming and read mapping statistics. See Table 1 in main text for explanations of datasets. | | | | | | | |
| --- | --- | --- | --- | --- | --- | --- | --- |
| Dataset | Species | Percent of Q30 bases before trimming | Percent of Q30 bases after trimming | Total reads before trimming | Percent of reads kept after trimming | Percent of trimmed reads mapping | Percent of mapped reads after removing duplicates |
| indSeq | DF | 88.92% | 90.94% | 419,786,724 | 98.07% | 85.48% | 68.68% |
|  | JP | 87.28% | 89.92% | 392,241,534 | 97.40% | 46.14% | 79.40% |
| poolSeq | DF | 88.76% | 90.44% | 109,897,366 | 99.05% | 86.31% | 41.55% |
|  | JP | 86.55% | 89.04% | 152,988,120 | 98.06% | 47.21% | 70.24% |
| megaSeq | DF | 95.63% | 97.55% | 203,296,026 | 97.16% | 83.52% | 99.65% |
|  | JP | 74.82% | 78.69% | 201,317,556 | 96.49% | 12.72% | 36.70% |

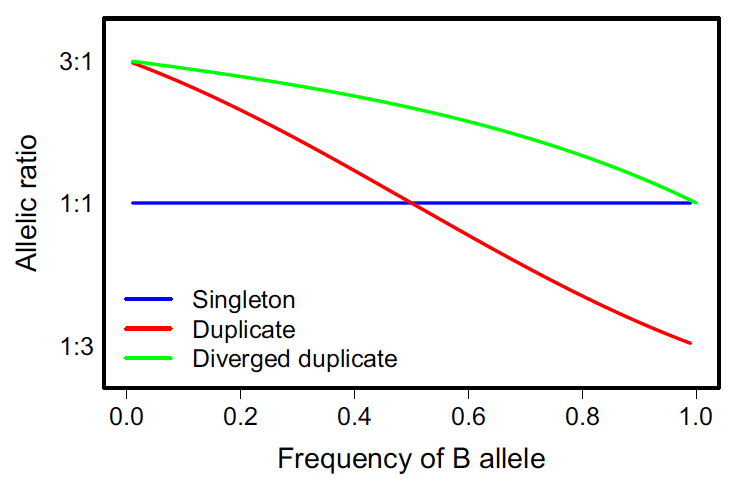

A
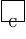

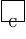

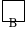

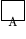

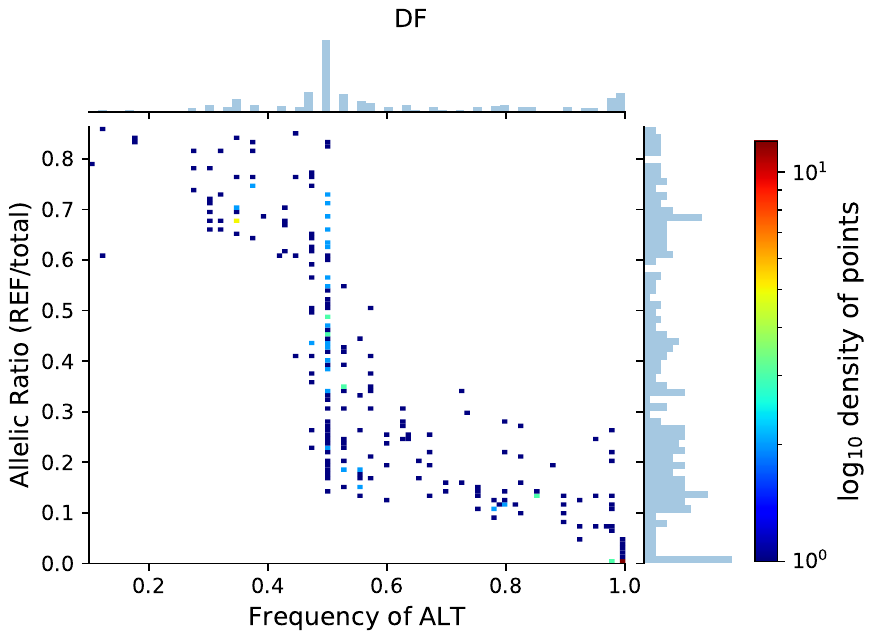

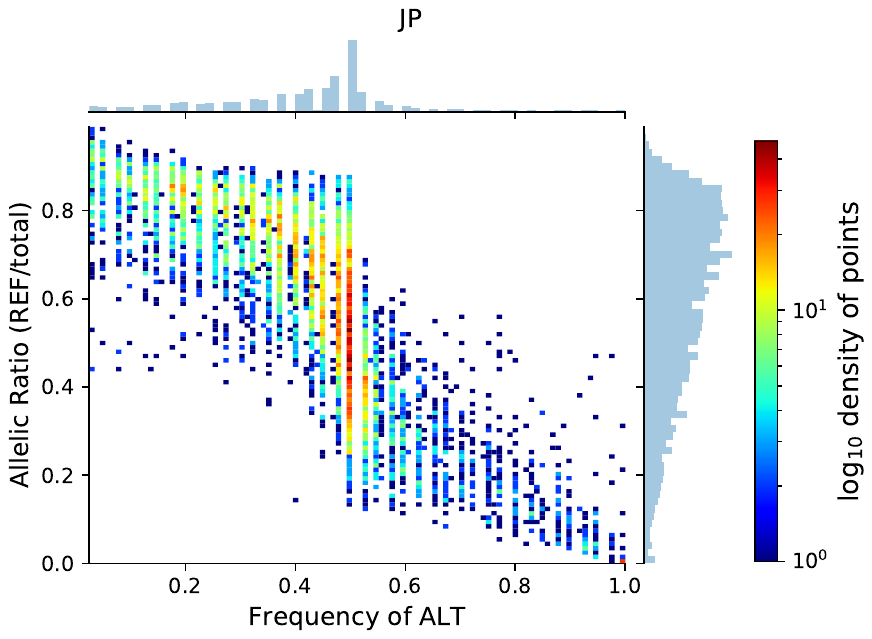

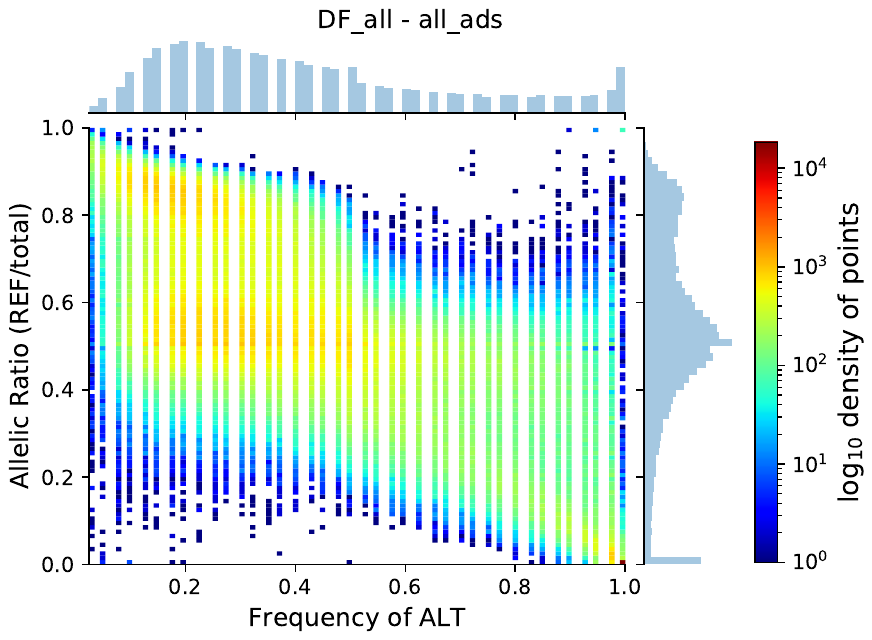

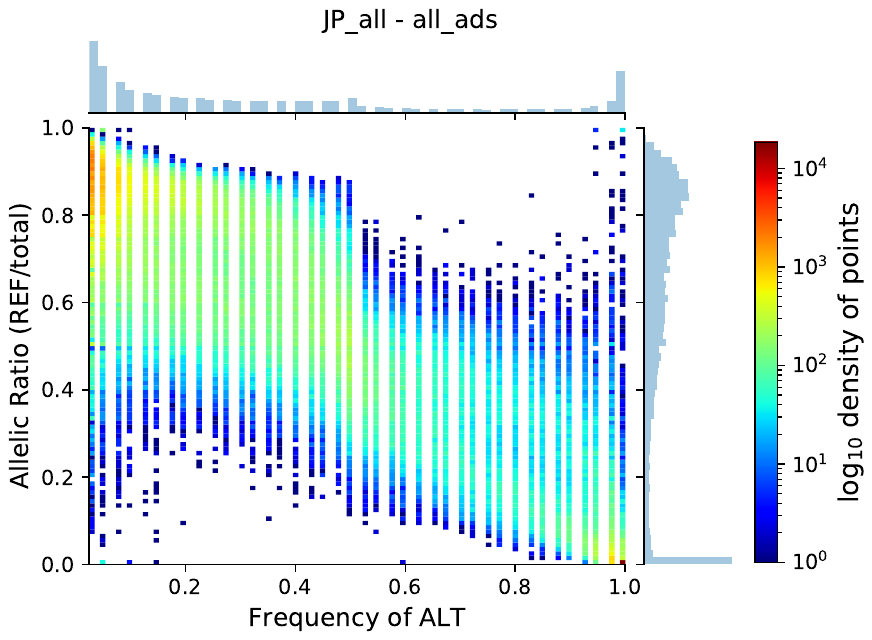

E

D

B

C

**Figure S1. Allelic ratio (the ratio of reference, REF, reads to the total number of reads) and the frequency of the alternative (ALT) allele calculated from indSeq data distinguish paralog misalignments according to expectations (A, Fig. 2 from McKinney et al. 2017), and empirically for Douglas-fir (B, D) and jack pine (C, E). B-C) Empirical distribution of megaSNP sites (candidate paralog sites identified as heterozygote calls from haploid tissue) calculated using indSeq data for those sites that were also called in poolSeq data (i.e, intersection I1). D-E) Empirical distribution of intersection I2 (indSeq and poolSeq intersection) calculated using indSeq data. Note color scale changes for each figure.**

A
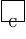

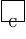

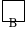

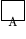

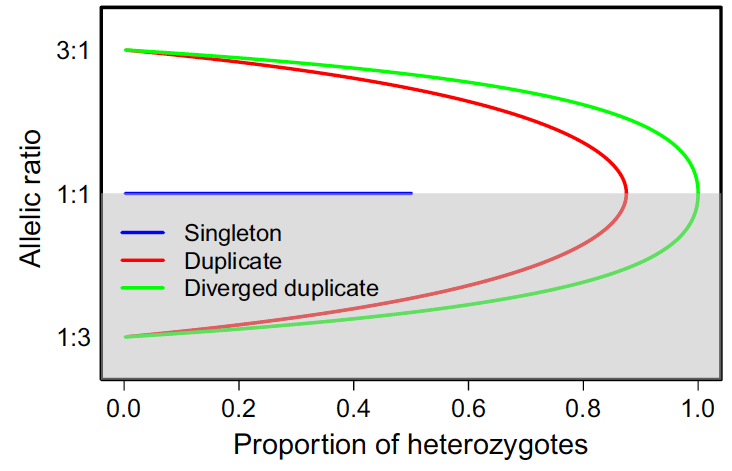

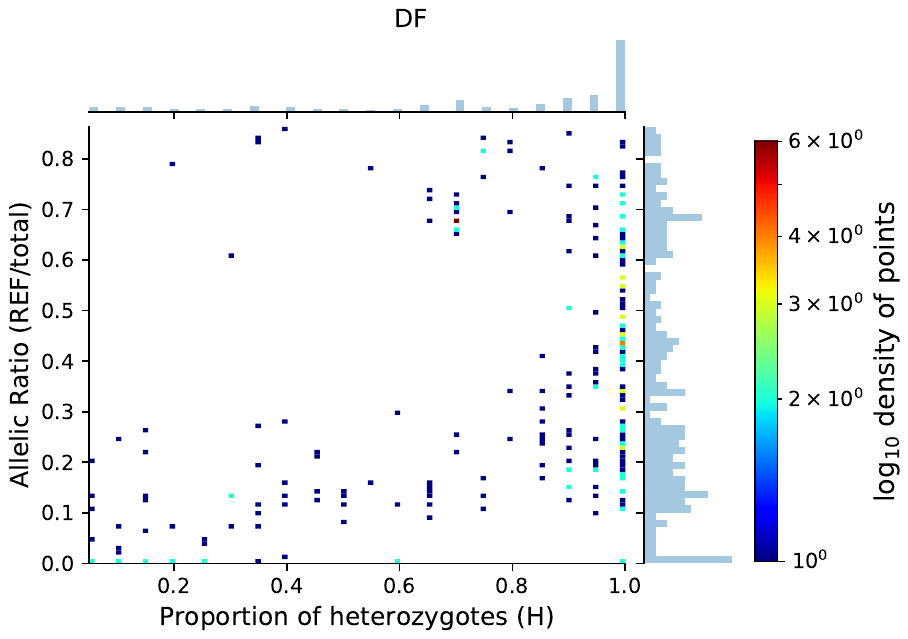

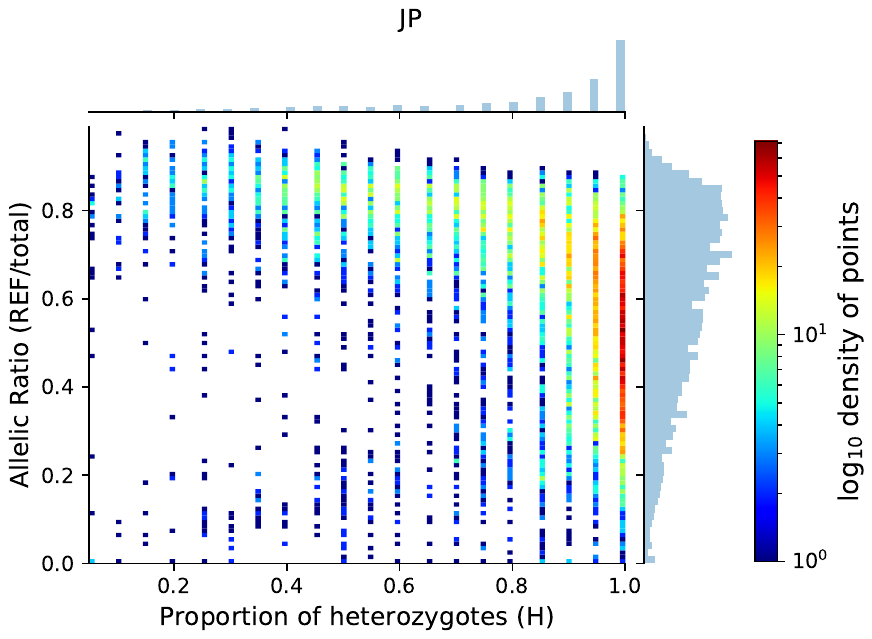

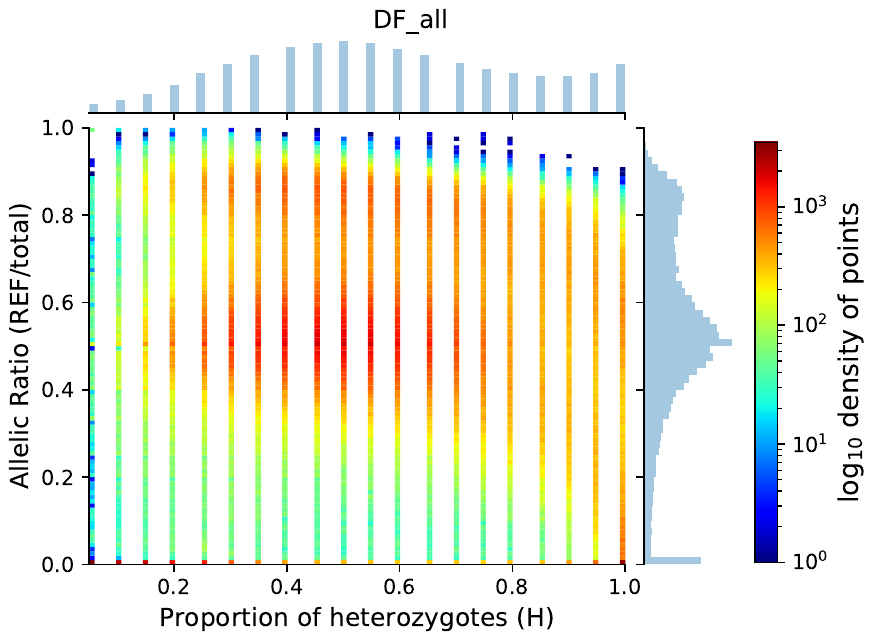

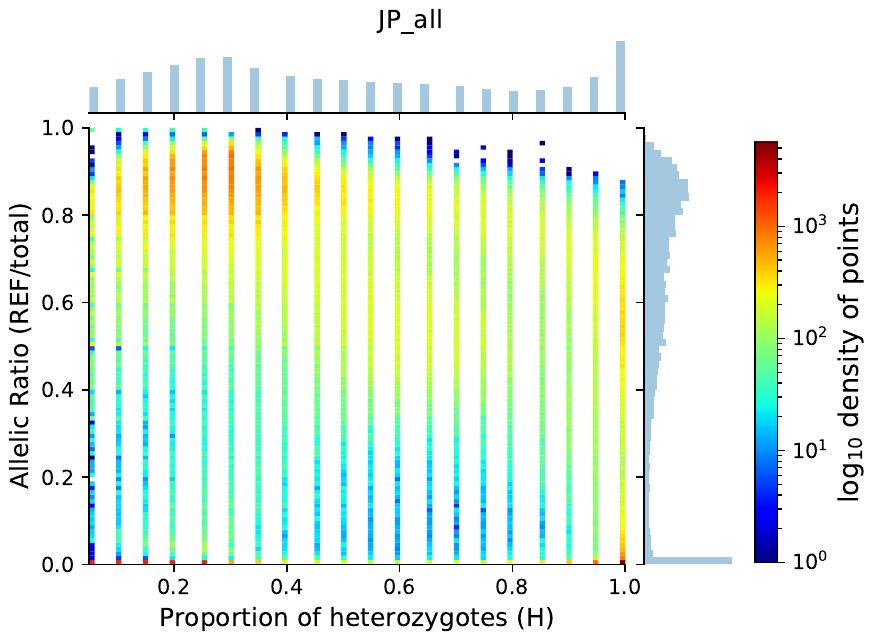

B

C

D

E

**Figure S2. Allelic ratio (the ratio of reference, REF, reads to the total number of reads) and the proportion of heterozygotes, *H*, calculated from indSeq data distinguish paralog misalignments according to expectations (A, Fig. 3 from McKinney et al. 2017), and empirically for Douglas-fir (B, D) and jack pine (C, E). B-C) Empirical distribution of megaSNP sites (candidate paralog sites identified as heterozygote calls from haploid tissue) calculated using indSeq data for those sites that were also called in poolSeq data (i.e, intersection I1). D-E) Empirical distribution of intersection I2 (indSeq and poolSeq intersection) calculated using indSeq data. The grayed area in (A) is because this space is expected to be more apparent when randomly choosing the allele; we used the reference allele to calculate the allelic ratio. Note color scale changes for each figure.**

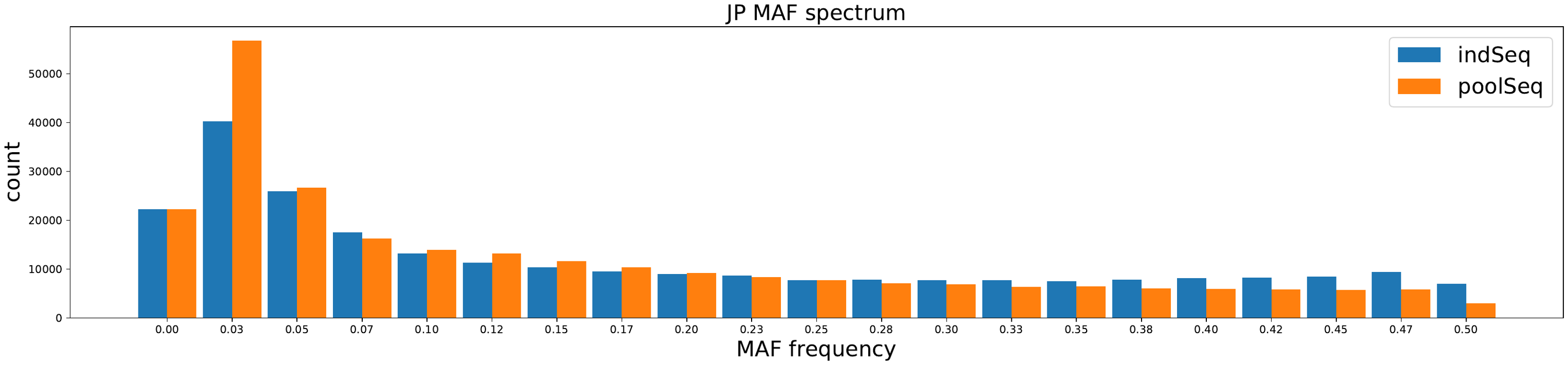

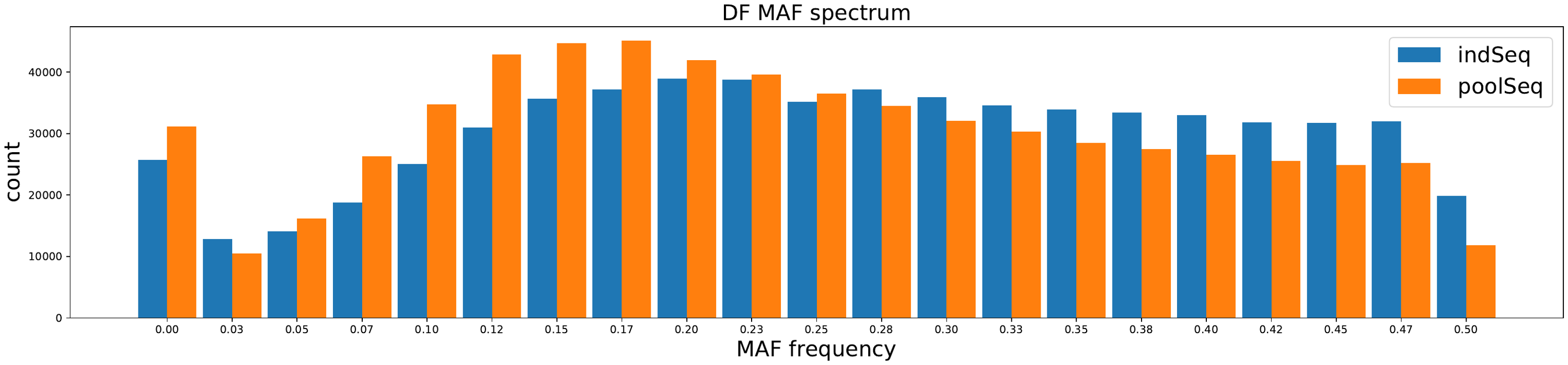

B

A
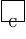

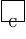

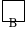

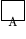

**Figure S3**. Comparison of the minor allele frequency spectra estimated for jack pine (A) and Douglas-fir (B) for indSeq and poolSeq datasets (see Table 1 of main text for descriptions of datasets).

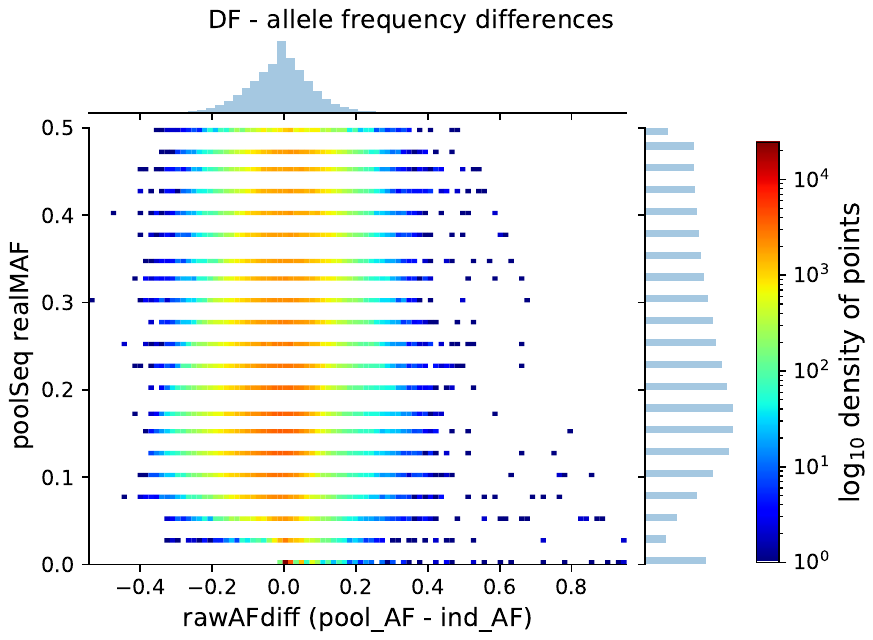

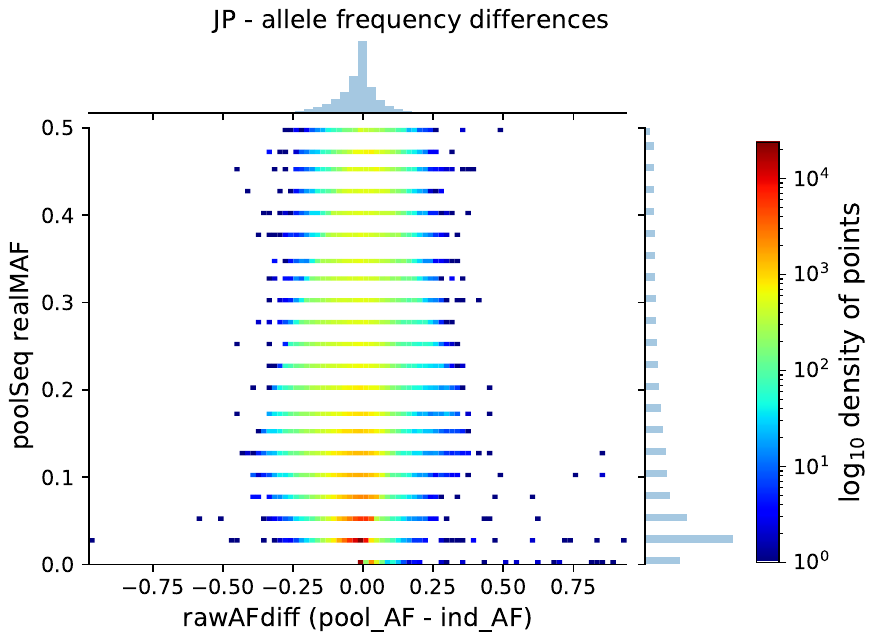

B

A
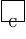

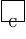

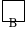

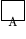

**Figure S4**. The difference between allele frequency estimates between poolSeq and indSeq datasets ($AFdiff$) against the poolSeq minor allele frequency estimated by VarScan for Douglas-fir (A) and jack pine (B). In general, there is little pattern between $AFdiff$ and poolSeq MAF (generally an approximately normally distributed about $AFdiff=0$), except for the lowest MAF bin.

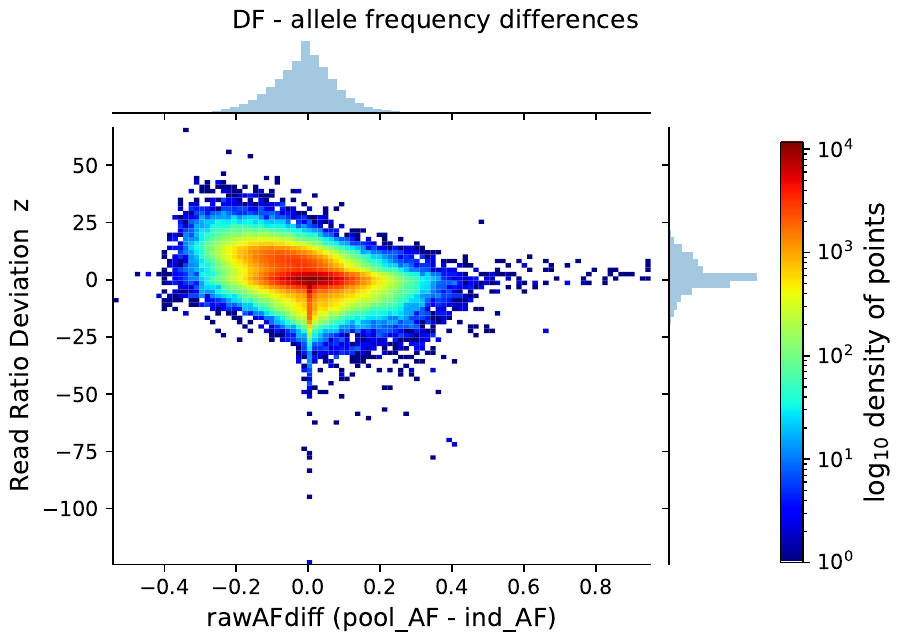

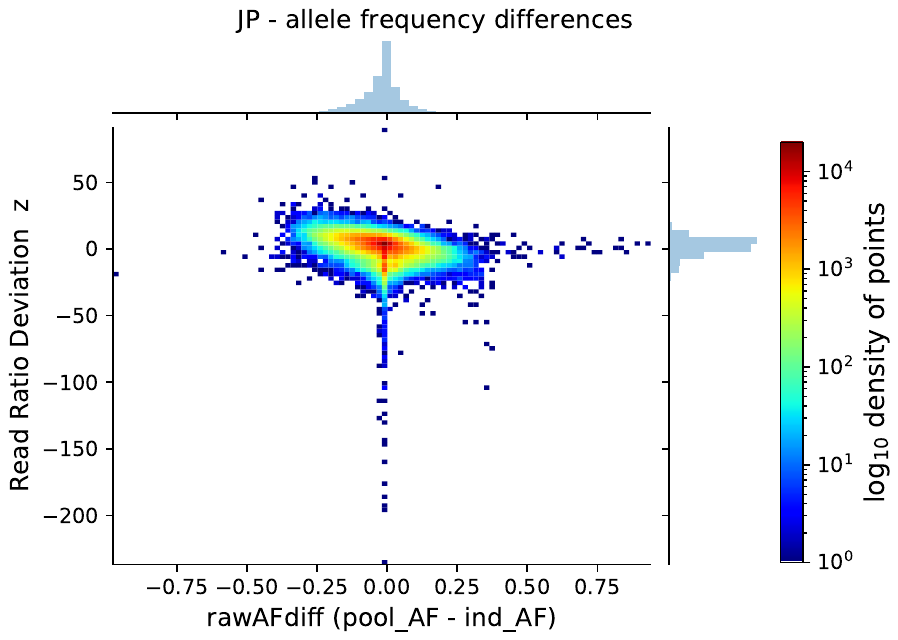

A
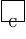

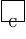

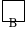

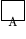

B

**Figure S5**. Differences in allele frequency estimates between datasets (AFdiff) is affected by the z-score of allele ratio deviations for Douglas-fir (A) and jack pine (B). Color scale is not standardized among figures to better accentuate patterns in our data.

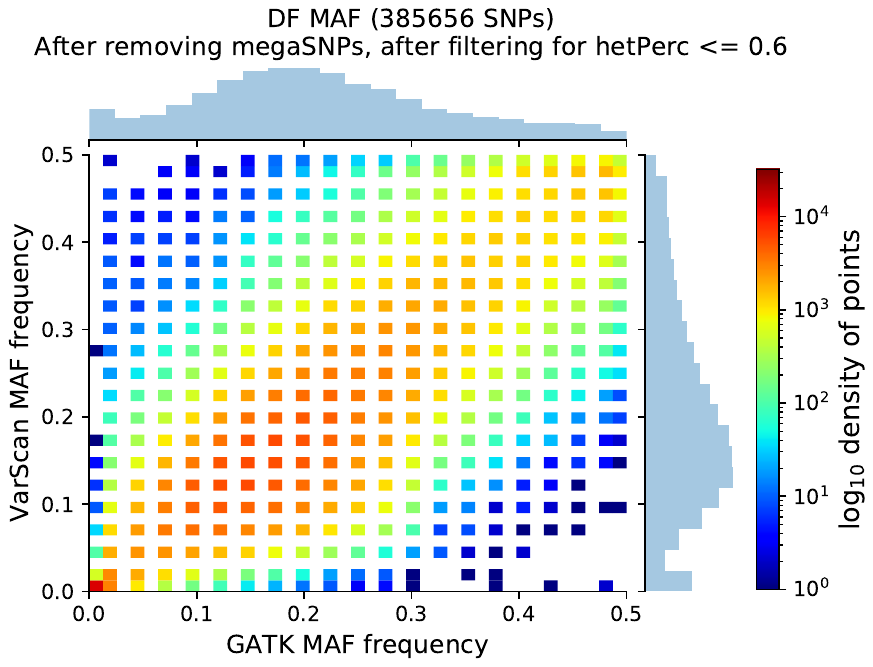

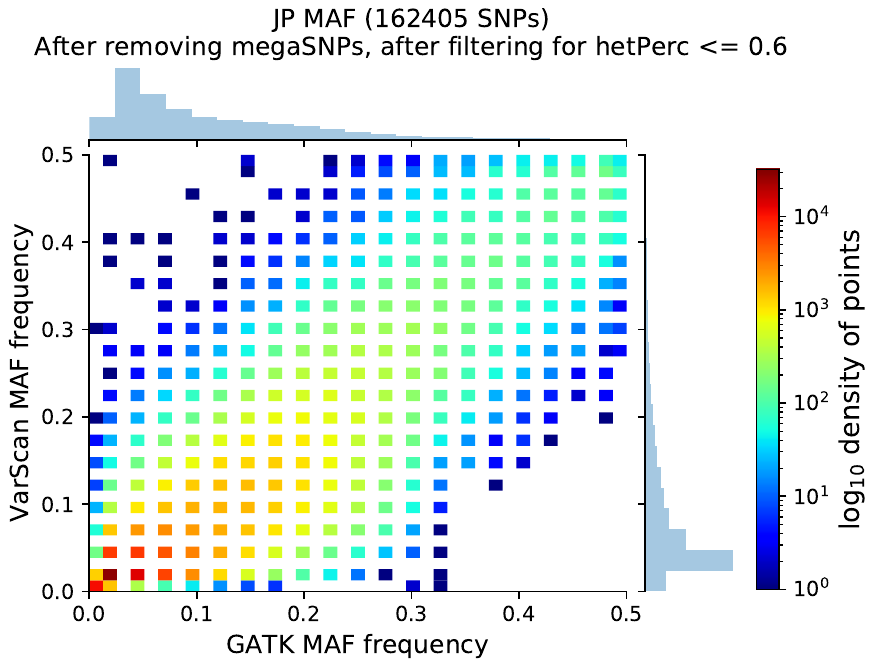

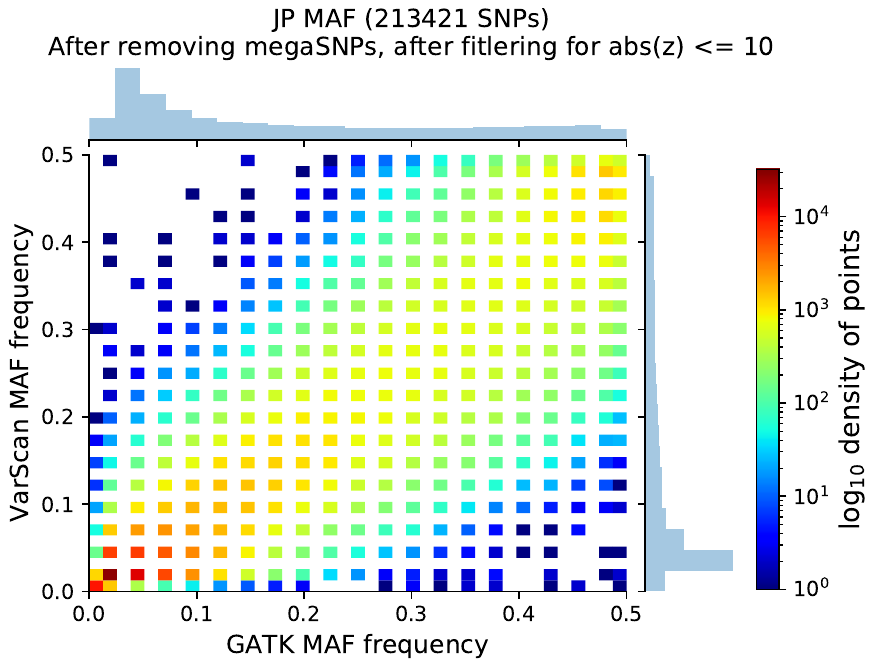

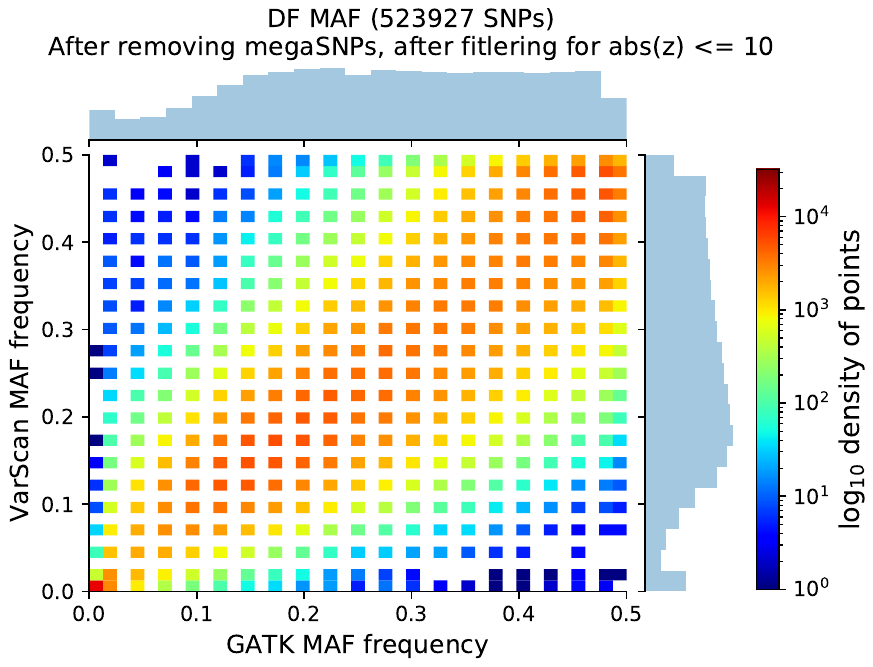

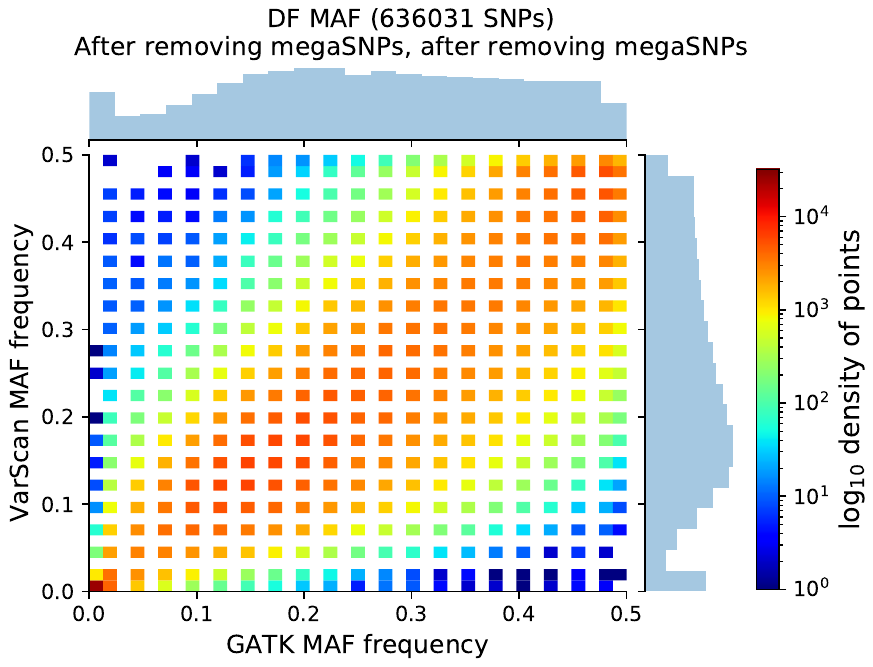

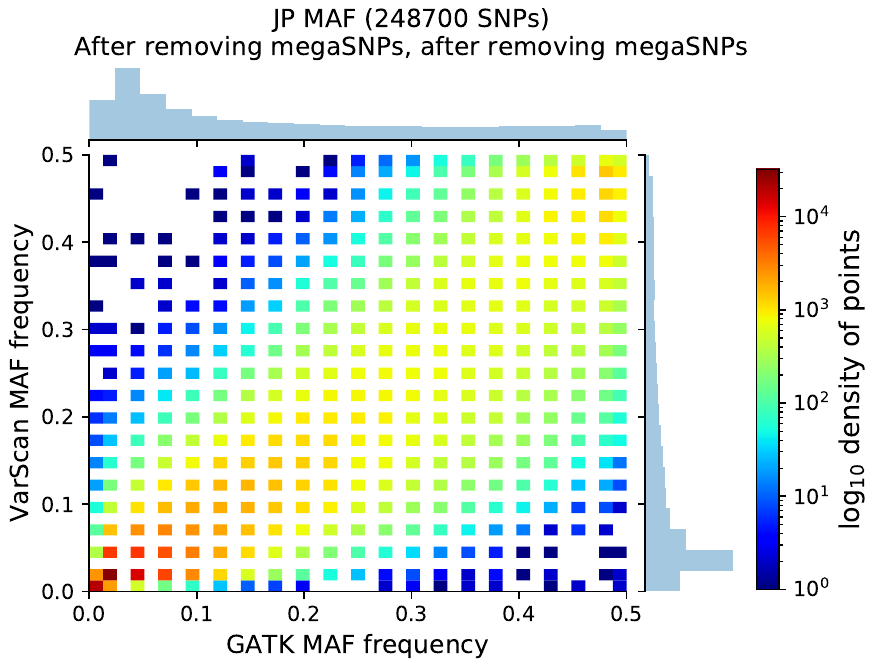

**Figure S6**. 2D histograms for Douglas-fir (A, C, E) and jack pine (B, D, F) displaying congruence between indSeq (x-axis) and poolSeq (y-axis) minor allele frequency estimates A-B) after removing candidate paralog sites (megaSNPs) and then subsequently C-D) removing sites with absolute $z$-score > 10; compare with Figure 5A-B of main text. Panels E-F show congruence between megaSNP-filtered data (A-B) after removing $H > 0.6$. Color scale is standardized across subfigures to visualize differences in density between filtering steps.

poolSeq MAF frequency

poolSeq MAF frequency

indSeq MAF frequency

indSeq MAF frequency

indSeq MAF frequency

indSeq MAF frequency

indSeq MAF frequency

indSeq MAF frequency

poolSeq MAF frequency

poolSeq MAF frequency

poolSeq MAF frequency

poolSeq MAF frequency

F

E

B

A
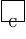

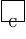

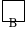

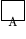

D

C

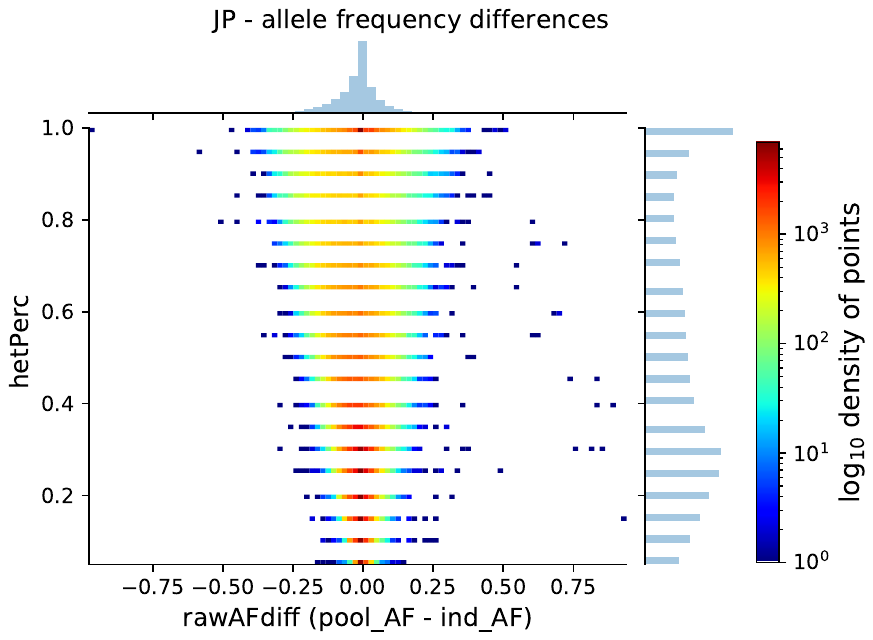

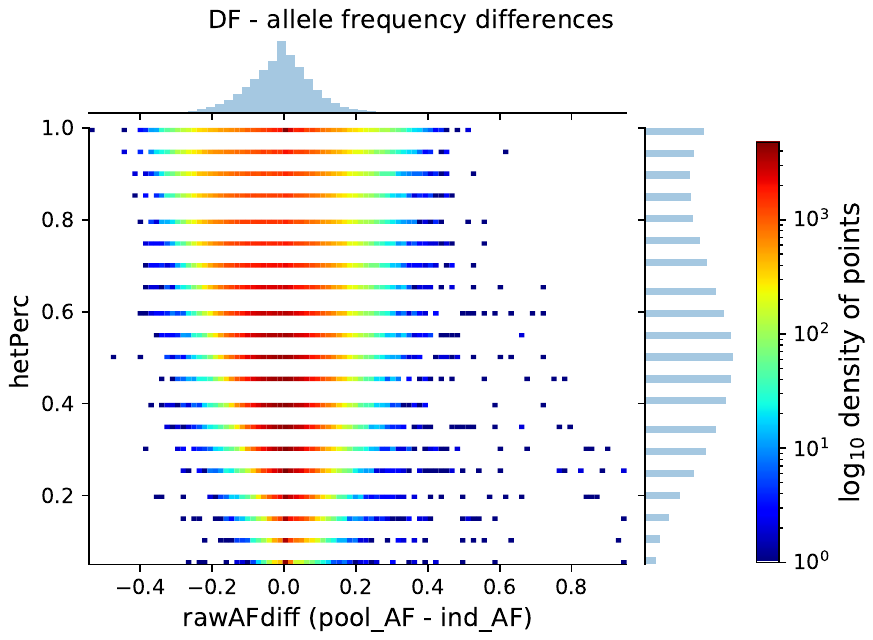

B

A
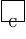

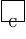

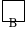

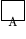

**Figure S7**. Difference in allele frequency estimates between poolSeq and indSeq ($AFdiff$) as influenced by the percentage of heterozygotes found at indSeq sites ($H$, y-axis). This intersection represents filtering intersection I2 for megaSNP sites.

### 3 | Supplemental References

García C, G Escribano-Ávila (2016) An optimised protocol to isolate high-quality genomic DNA from seed tissues streamlines the workflow to obtain direct estimates of seed dispersal distances in gymnosperms. Journal of Plant Research 129, 559–563.

[dataset] Neale DB, PE McGuire, NC Wheeler, KA Stevens, MW Crepeau, C Cardeno, AV Zimin, D Puiu, GM Pertea, UU Sezen, C Casola, TE Koralewski, R Paul, D Gonzalez-Ibeas, S Zaman, R Cronn, M Yandell, C Holt, CH Langley, JA Yorke, SL Salzberg, JL Wegrzyn (2017c) Plant sample from *Pseudotsuga menziesii*. Sequence Read Archive of the National Center for Biotechnology Information (NCBI SRA). Version 1.0. BioSample: SAMN03333061; Sample name: Psme_Weyco1; SRA: SRS937347.
